# NELL-1 regulates the matrisome to alter osteosarcoma disease progression

**DOI:** 10.1101/2022.01.21.477245

**Authors:** Qizhi Qin, Mario Gomez-Salazar, Robert J. Tower, Leslie Chang, Carol D. Morris, Edward F. McCarthy, Kang Ting, Xinli Zhang, Aaron W. James

## Abstract

Sarcomas produce abnormal extracellular matrix (ECM) which in turn provides instructive cues for cell growth and invasion. Neural EGF Like-Like molecule 1 (NELL1) is a secreted glycoprotein characterized by its non-neoplastic osteoinductive effects, yet highly expressed in skeletal sarcomas. Here, *NELL1* gene deletion markedly reduced invasive behavior across human osteosarcoma (OS) cell lines. This resulted in reduced OS disease progression, inhibited metastatic potential and improved survival in a xenograft model. These observations were recapitulated with *Nell1* conditional knockout in mouse models of p53/Rb driven sarcomagenesis, including reduced tumor frequency, and extended tumor free survival.

Transcriptomic and phospho-proteomic analysis demonstrated that *NELL1* loss skews the expression of matricellular proteins associated with reduced FAK signaling. Culture on OS enriched matricellular proteins reversed phenotypic and signaling changes among *NELL1* knockout sarcoma cells. These findings in mouse and human models suggest that NELL1 expression alters the sarcoma matrix, thereby modulating cellular invasive potential and prognosis. Disruption of NELL1 signaling may represent a novel therapeutic approach to short circuit sarcoma disease progression.

## Introduction

Sarcomas are a broad group of malignant neoplasms that arise from mesenchymal cells origin in soft tissue and bone ^1, 2^. Emerging molecular findings suggest that mechanical and chemical properties of the tumor microenvironment act together to accelerate sarcoma disease progression^3^. Within the tumor microenvironment, structural extracellular matrix (ECM) proteins play crucial role in cell surface receptor signaling ^4, 5^, which in turn may regulate sarcoma cell invasive and metastatic potential ^6–10^. Osteosarcoma (OS), the most common primary malignancy of bone ^11–14^, is defined by its pathogenic osseous ECM ^15^. Several studies have demonstrated OS matrix is extensively altered, including changes in collagens and proteoglycans ^15, 16^. In particular, high expression of several sarcomatous matrix proteins has been associated with poor response to chemotherapy and poor prognosis in clinical studies of OS ^17–20^.

NELL1 (Neural EGFL like 1) is a secreted osteoinductive protein which has bone anabolic ^21–24^ and anti-osteoclastic effects ^25, 26^. NELL1 can significantly stimulate mesenchymal stromal cell proliferation and osteogenic differentiation, until recently NELL1 was simply known to regulate skeletal patterning and induce bone defect healing in small and large animals ^27–29^. Earlier studies revealed that loss of *Nell1* expression or function in mice reduces expression of numerous ECM proteins, as well as proteins involved in cell adhesion and cellular communication ^30^. Notably, NELL1 expression has been detected in bone tumors and cartilage-forming tumors ^31^. With these recent insights, it is likely that the cellular effects of NELL1 are much more pleiotropic than once understood. Nevertheless the biologic implications of NELL1 signaling in osteosarcomagenesis and/or disease progression have been until this point unknown. Here, we sought to define the regulatory role of *NELL1* in sarcoma pathogenesis. CRISPR/Cas9 mediated *NELL1* gene deletion markedly reduced the malignant phenotype across human OS cell lines. *NELL1* knockout significantly slowed OS disease progression, blunted metastatic potential, and improved survival in a xenograft model. Similar observations were identified with *Nell1* gene deletion in a p53/Rb driven osteosarcomagenesis mouse model. Transcriptomic and proteomic analysis demonstrated *NELL1* knockout skews the expression of key matricellular proteins associated with reduced FAK signaling. These findings in mouse and human osteosarcoma suggest that NELL1 signaling positively regulates multiple aspects of OS disease progression in part via alterations in the sarcomatous ECM, and that disruption of NELL1 signaling may represent a novel therapeutic approach.

## Results

### Cell autonomous effects of NELL1 gene deletion in osteosarcoma

First, expression of *NELL1* transcripts was observed to be a conserved feature across all human OS cell lines examined, a finding in agreement with prior observations (**Fig. 1A**) ^31^. Gene deletion studies were first performed using the 143B cell line, generating six *NELL1* KO clones using CRISPR/Cas9. Knockout was confirmed using qRT-PCR, T7 endonuclease I assay (**Supplementary Fig. S1**) and immunocytochemistry (**Fig. 1B**). In comparison to vector control, multiple cellular effects were observed among *NELL1* KO cell clones, including reduced proliferation (**Fig. 1C**, mean 34.9% reduction at 72 h), attachment (**Fig. 1D, E**, mean 31.9% reduction at 5 h), and invasion (**Fig. 1F, G**, 25.1-69.1% reduction across clones). Similar findings were observed with analysis of a polyclonal *NELL1* KO 143B cell population (**Supplementary Fig. S2**), and indeed similar assessments across five other available OS cell lines recapitulated these findings (**Fig. 1H-J**). Overexpression studies were performed, in which adenoviral delivery (Ad-*NELL1*) led to partial restoration of gene expression and near complete restoration of attachment and invasion potential among *NELL1* KO 143B sarcoma cells (**Fig. 1K-M**). These data confirmed that the *NELL1* gene plays a crucial role in maintaining cellular proliferation, attachment, and invasion potential in OS cells.

**Figure 1.**
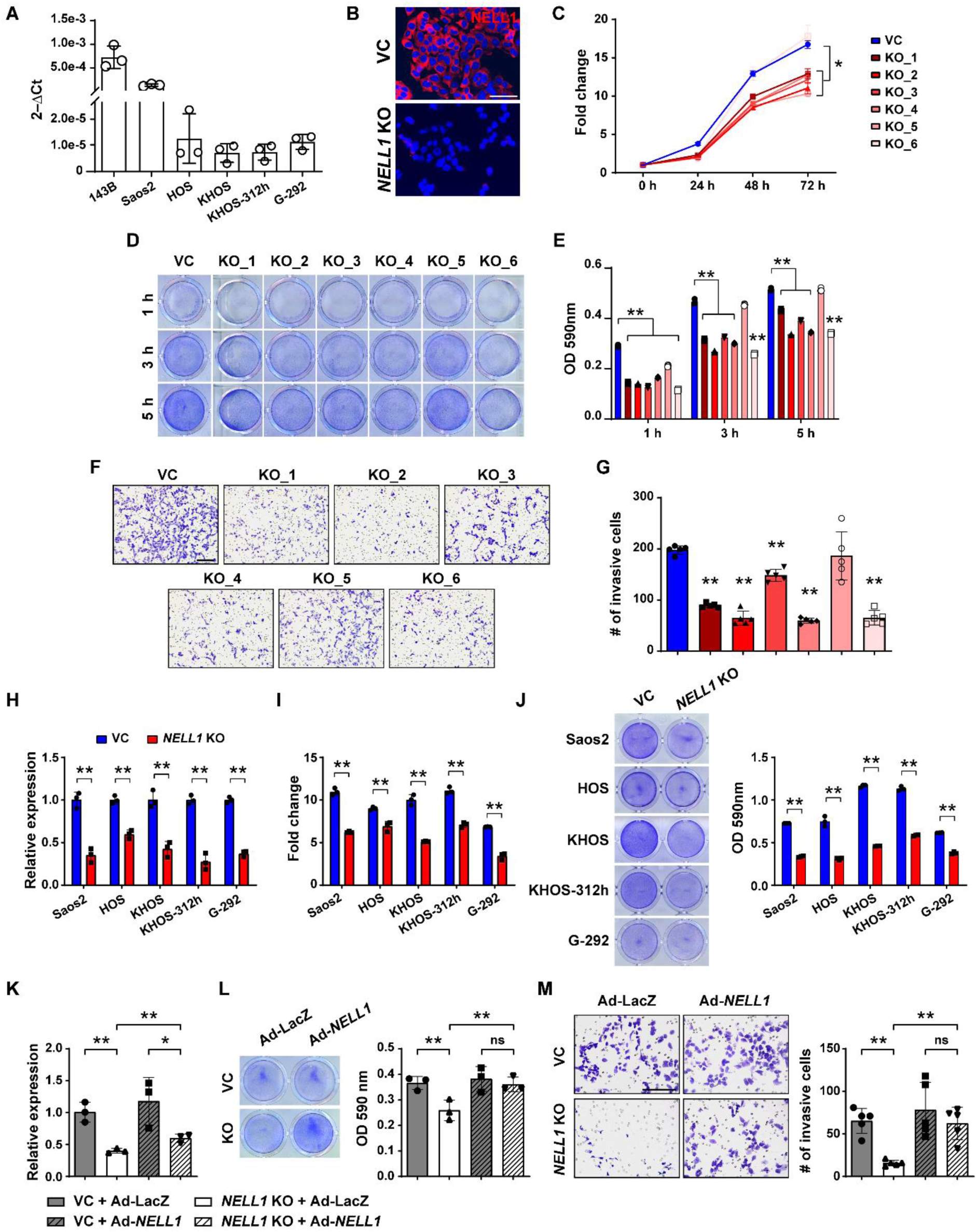
Cell autonomous effects of CRISPR/Cas9-mediated *NELL1* gene deletion in human osteosarcoma cell lines. **(A)** *NELL1* expression by qRT-PCR in osteosarcoma cell lines. **(B-G)** Effects of *NELL1* gene deletion in 143B cells. **(B)** Confirmation of *NELL1* knockout (KO) in single cell clones by representative immunocytochemical staining, in comparison to vector control (VC). Additional data shown in **Supplementary Figure S1**. **(C)** MTS proliferation assay among *NELL1* KO single cell clones at 24, 48, and 72 h. **(D, E)** Attachment assay at 3 and 5h assessed by (**D**) crystal violet staining and (**E**) quantification. **(F, G)** Transwell invasion assay at 24 h, including (**F**) representative images and (**G**) quantification. (**H-J**) Effects of CRISPR-mediated *NELL1* gene deletion across other human OS cell lines, including Saos2, HOS, KHOS, KHOS312h, and G292 cells. **(H)** Confirmation of *NELL1* KO by qRT-PCR. **(I)** MTS proliferation assay at 72 h. (**J**) Attachment by crystal violet staining and quantification at 5 h. (**K-M**) Effects of adenoviral *NELL1* (Ad-*NELL1*) among 143B cells with or without *NELL1* gene deletion. Experiments performed in comparison to Ad-LacZ control. (**K**) *NELL1* gene expression by qRT-PCR. (**L**) Attachment assay by crystal violet staining and quantification at 5 h. (**M**) Transwell invasion assay and quantification at 24 h. Data shown as mean ± 1 SD, with dots representing individual datapoints. All experiments performed in triplicate replicates, with results from a single replicate shown. \**P*<0.05; \*\**P*<0.01; ns: non-significant. Scale bars: 500 µm. A two-tail Student’s *t* test was used for two group comparisons (Fig. H, I, J), and one-way ANOVA with post hoc Tukey test when more than two groups were compared (Fig. C, E, G, K, L, M).

**Figure 2.**
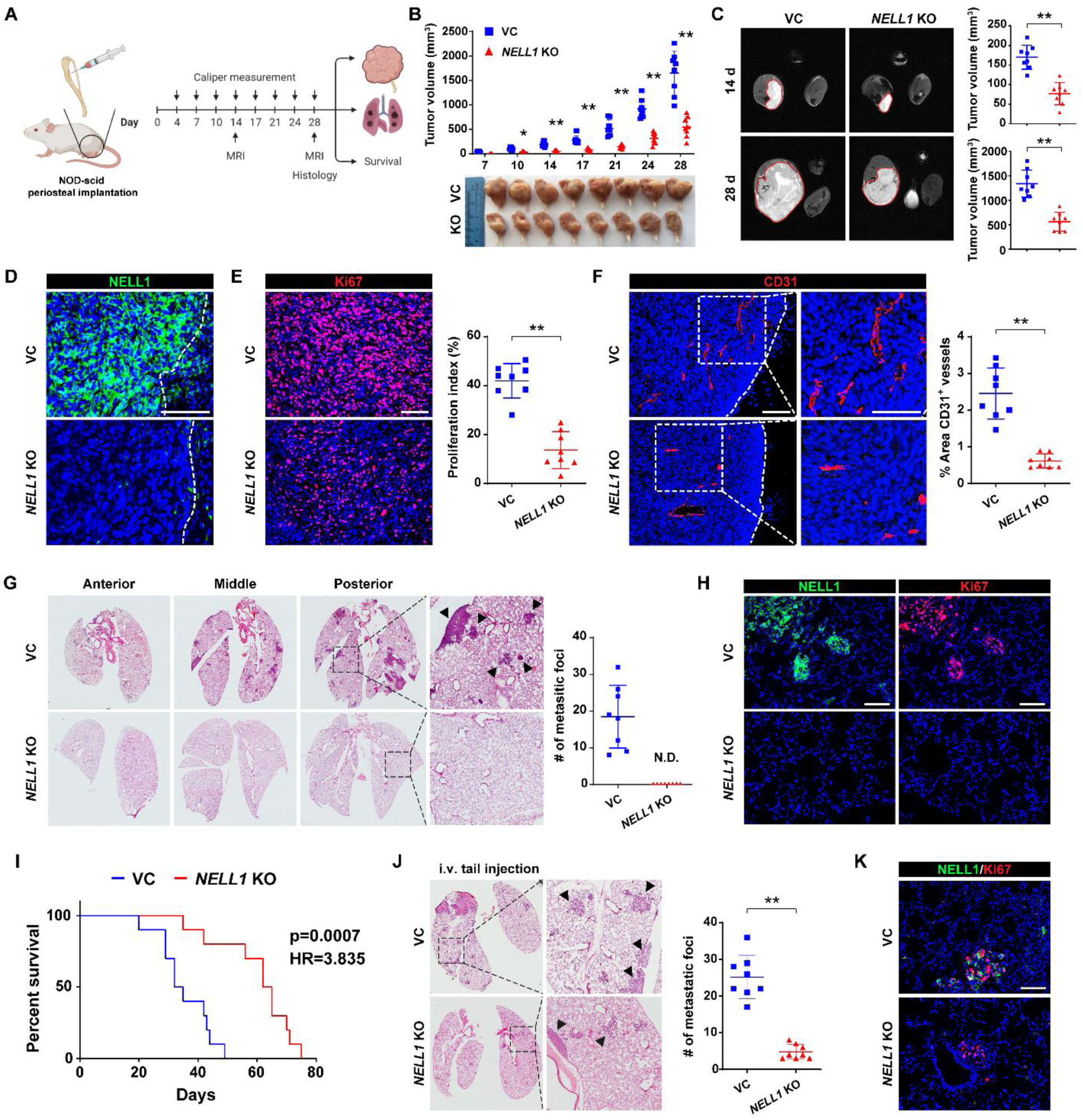
*NELL1* knockout mitigates OS disease progression in 143B xenograft model. Orthotopic implantation of *NELL1* KO or vector control clonal 143B cells within the proximal tibia of female NOD-Scid mice (n=8 mice per group, 1 × 10^6^ per implant). Biological replicate shown in **Supplementary Fig. S4**. **(A)** Schematic diagram of study design. **(B)** Tumor volume, calculated by caliper measurements twice weekly until 28 d post-injection (above) and gross pathology of all tumors (below). **(C)** Representative MRI imaging at 14 and 28 d post-injection (left) and tumor volume (right). **(D)** Confirmation of *NELL1* KO in xenograft tumors by immunostaining. Dashed line indicates edge of tumor. **(E)** Ki67 immunostaining (left) and quantification (right). **(F)** CD31 immunostaining (left) and quantification (right). Dashed line indicates edge of tumor. **(G)** Cross-sections of pulmonary fields (left) and quantification of metastatic foci (right). **(H)** Representative NELL1 and Ki67 immunostaining in lung metastatic foci. **(I)** Overall survival, evaluated using Kaplan–Meier curves (n=10 mice per group, 1 × 10^6^ per implant). **(J)** *NELL1* gene deletion mitigates OS lung metastasis after systemic injection. Tail vein injection of *NELL1* KO or vector control 143B cells in female NOD scid mice, followed by analysis after 28 d (1.5 × 10^6^ cells / mouse, n=8 mice per group). Cross-sections of pulmonary fields (left) and quantification of metastatic foci (right). **(K)** Representative NELL1 and Ki67 immunostaining in lung metastatic foci. Scale bar: 100 µm. Individual dots in scatterplots represent values from single animal measurements, while mean and one SD are indicated by crosshairs and whiskers. \**P*<0.05; \*\**P*<0.01. ND: Not detected. A two-tailed student’s *t* test was used for all comparisons. Survival was assessed using Kaplan–Meier curves and log-rank analysis (Fig. I).

### NELL1 knockout mitigates OS disease progression

*NELL1* KO 143B clones were next implanted in an orthotopic xenograft model in NOD scid mice (**Fig. 2A**). In relation to control, tumor growth was impeded as assessed by serial caliper measurements (**Fig. 2B**, 66.9% reduction in tumor size at study endpoint). Findings were confirmed in an independent clone of knockout tumor cells (**Supplementary Fig. S3**, 89.2% reduction in tumor size). MRI confirmed a 57.1% and 66.1% reduction in tumor size at 14 and 28 d post-implantation (**Fig. 2C**). Histologic examination of primary tumors demonstrated complete loss of NELL1 immunoreactivity among *NELL1* KO xenograft tumor cells (**Fig. 2D**), accompanied by a reduction in the proliferative index of *NELL1* KO implants (**Fig. 2E**, 67.5% reduction in Ki67 labeling). Tumor associated angiogenesis also showed a notable reduction among *NELL1* KO implants (**Fig. 2F**, 74.9% reduction in CD31-positive vessel area), in line with prior reports of the angiogenic effects of NELL1 signaling ^32–34^. In order to confirm the impact of NELL1 on invasion and metastasis, pulmonary metastatic burden was assessed by serial sections of lung fields. Results indicated complete absence of lung metastasis among *NELL1* KO tumors at 28 d post-inoculation (**Fig. 2G, H**). In a separate cohort of animals, mice with *NELL1* KO tumors demonstrated prolonged overall survival (**Fig. 2I**, 89.6% increase in median overall survival). Reduction in metastatic potential among *NELL1* KO sarcoma cells was further verified using tail vein injection, in which an 81.1% reduction in pulmonary burden was observed (**Fig. 2J, K**). Together, our data demonstrates that *NELL1* gene deletion slows OS tumor growth and mitigates metastasis in a xenograft model.

### NELL1 gene deletion impedes osteosarcomagenesis

The consequences of *NELL1* gene deletion were next examined in two murine OS models ^35, 36^. Firstly, mouse BMSCs were isolated from *p53*^fl/fl^;*Rb*^fl/fl^ (DKO) or *p53*^fl/fl^;*Rb*^fl/fl^;*Nell1*^fl/fl^ (TKO) mice, recombination performed via adenovirus-mediated Cre delivery and the consequences examined *in vitro* and after periosteal transplantation (**Fig. 3A**). As expected, gene knockout of *p53* and *Rb* resulted in an increased proliferative and invasive phenotype in comparison to vector control (**Supplementary Fig. S4**). In comparison to DKO tumor cells, TKO cells demonstrated reduced colony formation (**Fig. 3B**, 33.3% reduction), reduced cellular attachment (**Fig. 3C**, 27.5% and 19.8% reduction at 3 and 5 h, respectively), and reduced tumorsphere formation efficiency (**Fig. 3D**, 35.9% reduction at 5 d). DKO and TKO cells were next orthotopically implanted in NOD scid mice, and sarcomagenesis and growth were monitored (**Fig. 3E-H**). Tumor formation rates were significantly reduced among TKO implants (25.0% incidence in comparison to 100% incidence among DKO implants, **Supplementary Table S1**). Among those implants with tumor formation, tumor size was significantly lower in TKO cell implants (**Fig. 3E**, 76.7% reduction in tumor size). Histological analysis confirmed an osteosarcoma-like appearance of all tumors, and confirmed complete loss of NELL1 immunoreactivity among TKO tumor cells (**Fig. 3F**). In similarity to prior findings, *Nell1* gene deletion demonstrated a reduction in proliferative index and vascularity among tumors, assessed by Ki67 and CD31 immunostaining (**Fig. 3G**, 86.9% reduction in Ki67, and 71.1% reduction in CD31 immunostaining). Moreover, tumor-free survival was significantly prolonged in mice with TKO implants, with a hazard ratio (HR) of 0.115 (**Fig. 3H**).

**Figure 3.**
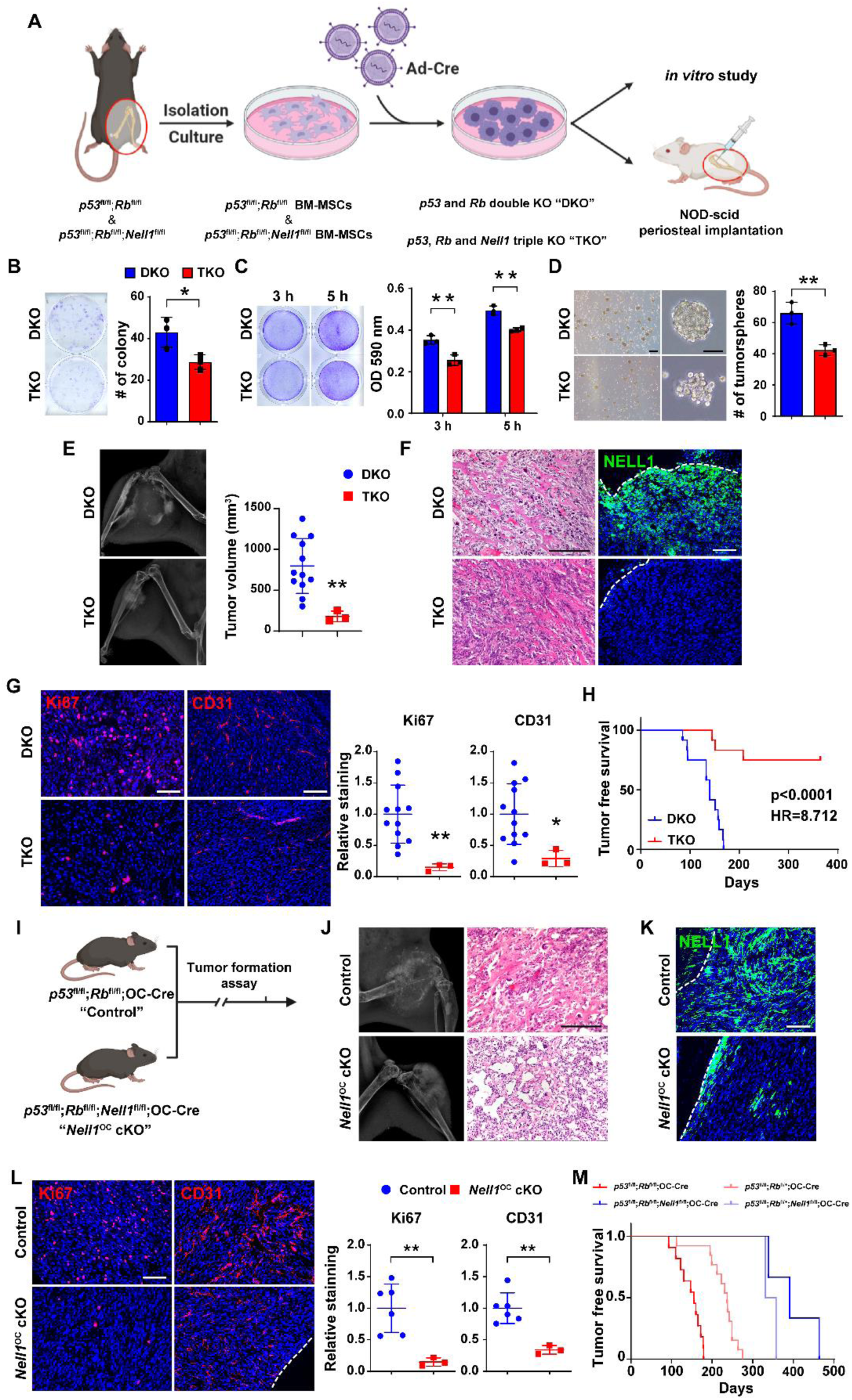
*Nell1* gene deletion slows osteosarcomagenesis in mouse lines. **(A)** Schematic diagram of study design. **(B-D)** Effects of Adeno-Cre treatment in *p53*^fl/fl^;*Rb*^fl/fl^ (DKO) or *p53*^fl/fl^;*Rb*^fl/fl^;*Nell1*^fl/fl^ (TKO) mouse bone marrow mesenchymal stromal cells. **(B)** Colony-forming unit (CFU) assay, assessed by crystal violet staining (left) and quantification (right) at 10 d. **(C)** Attachment assay assessed by crystal violet staining (left) and quantification (right), at 3 and 5 h. **(D)** Tumorsphere formation assay images (left) and quantification (right) at 5 d. **(E)** Osteosarcomagenesis visualized by X-ray of representative tumors (left) and tumor volume calculated by caliper measurements (right). **(F)** Representative histologic appearance by routine H&E staining and NELL1 immunostaining among DKO and TKO tumor tissue. **(G)** Representative Ki67 and CD31 immunostaining (left) and quantification (right) among DKO and TKO tumor tissues. **(H)** Tumor-free survival among DKO and TKO implanted mice, evaluated using Kaplan–Meier curves (n=12 mice per group, 1 × 10^6^ per implant). **(I-M)** Effects of *Nell1* gene deletion in spontaneous osteosarcomagenesis. **(I)** Schematic diagram of spontaneous osteosarcomagenesis among control (*p53*^fl/fl^;*Rb*^fl/fl^;OC-Cre) and *Nell1*^OC^ cKO animals (*p53*^fl/fl^;*Rb*^fl/fl^;*Nell1*^fl/fl^;OC-Cre). **(J)** Representative X-ray images (left) and histologic appearance by routine H&E staining (right). **(K)** Representative NELL1 immunostaining among control and *Nell1*^OC^ cKO tumor tissue. **(L)** Representative Ki67 and CD31 immunostaining (left) and quantification (right) among control and *Nell1*^OC^ cKO tumors. **(M)** Tumor-free survival, evaluated using Kaplan–Meier curves. Scale bars: 100 µm. For (Fig. B-D), data shown as mean ± 1 SD with dots representing individual measurement, performed in triplicate experimental replicates. For (Fig. E, G, L), dots in scatterplots represent values from individual animal measurements, while mean and one SD are indicated by crosshairs and whiskers. \**P*<0.05; \*\**P*<0.01. A two-tailed student’s *t* test was used for all comparisons. For (Fig. H, M), survival was assessed using Kaplan–Meier curves and log-rank analysis.

**Figure 4.**
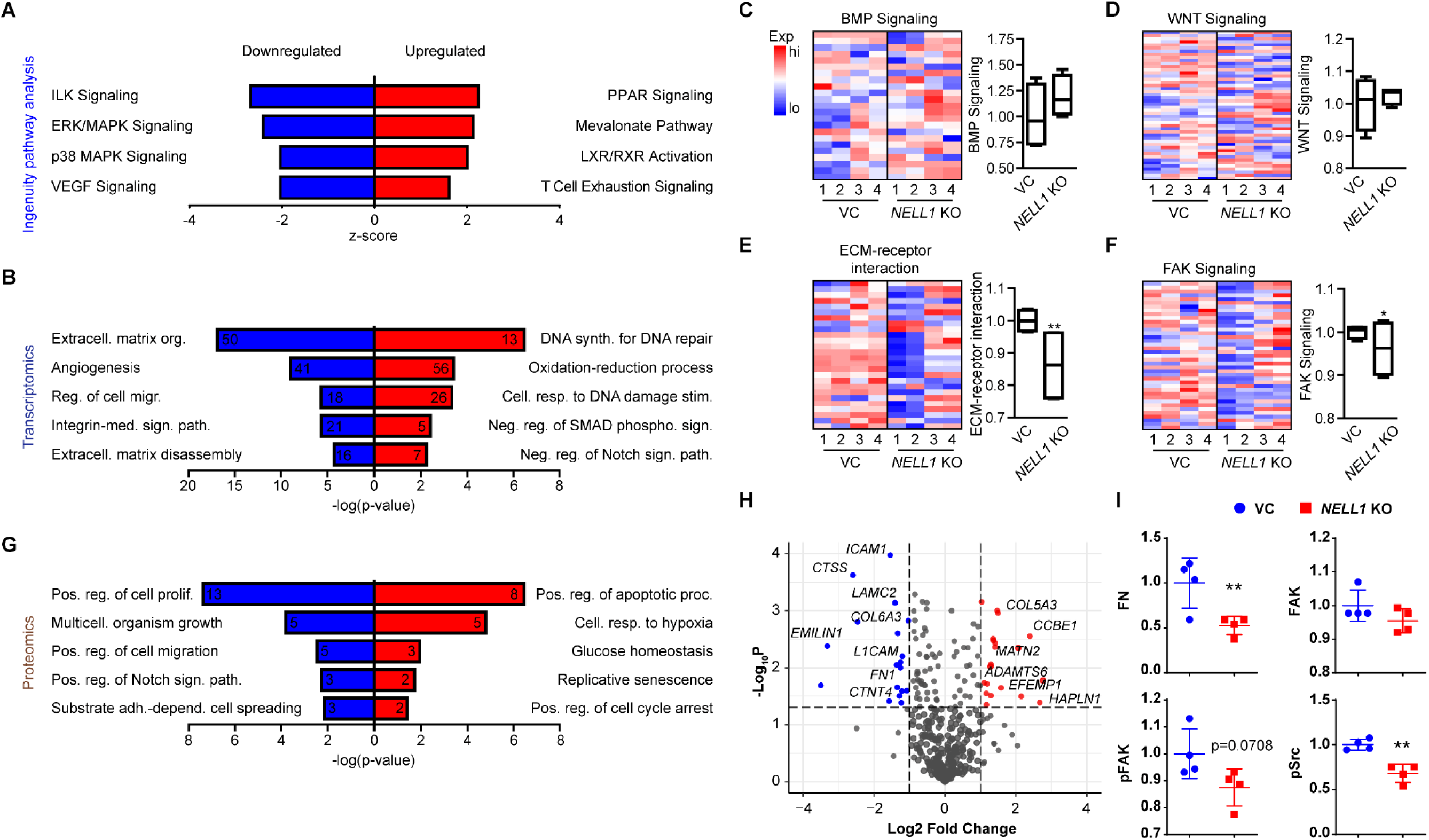
Bulk RNA sequencing and Reverse Phase Protein Arrays (RPPA) among *NELL1* KO 143B sarcoma cells. **(A)** Ingenuity Pathway Analysis (IPA) analysis of downregulated pathways (Z-score < 0; blue color) or upregulated pathways (Z-score > 0; red color) among *NELL1* KO cells in comparison to vector control (VC). **(B)** Gene Ontology (GO) enrichment analysis identified representative pathways that were downregulated (blue color) or upregulated (red color) among *NELL1* KO cells. **(C-F)** Heat map demonstrating expression levels of genes involved in *NELL1* downstream signaling and corresponding module score among *NELL1* KO as compared to VC cells, including **(C)** BMP signaling, **(D)** Wnt signaling, **(E)** ECM-receptor interaction, and **(F)** FAK signaling. **(G)** GO enrichment analysis identified representative pathways that were downregulated (blue color) or upregulated (red color) among *NELL1* KO cells as derived from RPPA. **(H)** Volcano plot demonstrating transcripts of ECM components that were downregulated (blue color) or upregulated (red color) in *NELL1* KO compared with VC genes derived from transcriptomics. **(I)** Protein expression of FAK signaling components derived from RPPA among *NELL1* KO cells. n=4 biological replicates of VC and *NELL1* KO cells. Dots in scatterplots represent values from individual measurement, while mean and one SD are indicated by crosshairs and whiskers. \**P*<0.05; \*\**P*<0.01. A two-tailed student’s *t* test was used for all comparisons.

Secondly, the same transgenes were used to monitor spontaneous osteosarcomagenesis, now utilizing the osteoblast specific Osteocalcin-Cre line ^37^. Animals with conditional knockout of *p53* and *Rb* (*p53*^fl/fl^;*Rb*^fl/fl^;OC-Cre, referred to as control) were compared to animals with conditional knockout of *p53, Rb* and *Nell1* (*p53*^fl/fl^;*Rb*^fl/fl^;*Nell1*^fl/fl^;OC-Cre, referred to as *Nell1*^OC^ cKO, **Fig. 3I**). Mice heterozygote for *Rb* were also examined for tumor incidence. Tumor formation rates were significantly reduced among *Nell1*^OC^ cKO animals (15.8% incidence in *Nell1*^OC^ cKO animals in comparison to 52.4% incidence among control animals, **Supplementary Table S2**). Radiographic and histologic appearance across tumors was consistent with osteosarcoma (**Fig. 3J**, tumor locations summarized in **Supplementary Table S3**). NELL1 immunohistochemical staining confirmed loss of NELL1 within sarcoma cells (**Fig. 3K**).

Immunohistochemical analyses again confirmed reduction in the proliferative index and tumor associated vasculature among tumors in *Nell1*^OC^ cKO animals (**Fig. 3L**, 85.2% reduction in Ki67, and 66.1% reduction in CD31 immunostaining). *Nell1*^OC^ cKO animals showed prolonged tumor-free survival (**Fig. 3M**, **Supplementary Table S3**, median tumor-free survival of 398 days in *Nell1*^OC^ cKO versus 146.5 days in controls). Collectively, *Nell1* loss impeded p53/Rb induced OS sarcomagenesis, but also reduced tumor growth and improved disease free survival in mouse models.

### NELL1 gene deletion shifts the sarcoma matrisome

Differences in gene expression profile were next investigated using bulk RNA sequencing across 4 clones of *NELL1* KO and VC 143B cells. Clear separation between gene expression profiles was observed between *NELL1* KO and control cells, as assessed by principal component analysis and volcano plot (**Supplementary Fig. S5A, B**). QIAGEN Ingenuity Pathway Analysis (IPA) showed that multiple signaling pathways were downregulated in *NELL1* KO cells, including for instance integrin-linked kinase (ILK), ERK/MAPK and VEGF signaling (**Fig. 4A**, see **Supplementary Table S4, 5** for a complete list). Next, pathway analysis of differentially expressed genes (DEGs) revealed Gene ontology (GO) term enrichment in extracellular matrix (ECM) organization, angiogenesis, integrin-mediated signaling pathway, and ECM disassembly were downregulated in *NELL1* KO cells (**Fig. 4B**, see **Supplementary Table S6, 7** for most downregulated and upregulated GO terms). Heat maps revealed differences in expression profile of OS related genes among *NELL1* KO and VC cells, including for example *MDM2, CDK4*, and *CD44* among others (**Supplementary Fig. S5C**) ^38–40^. Next, deletion of *NELL1* in OS cells was further validated by downstream signaling-related gene expression, shown by heatmap and cumulative module scoring (**Fig. 4C-F**). Although in other contexts canonical BMP and Wnt signaling pathway activation are known to be downstream of NELL1 signaling ^23, 41^, no significant change in either pathway was observed with *NELL1* gene deletion (**Fig. 4C, D**). Instead, significant reductions in gene modules related to ECM-receptor interaction and FAK signaling were observed among *NELL1* KO sarcoma cells (**Fig. 4E, F**).

In order to further evaluate differences with *NELL1* gene deletion, a reverse phase protein array was performed. A clear separation between protein profiles was observed when comparing *NELL1* KO and VC cells, as assessed by principal component analysis and volcano plot **(Supplementary Fig. S6A, B**). Again, pathway analysis of downregulated DEGs revealed GO term enrichment in *NELL1* KO cells associated with the regulation of cellular proliferation, migration, adhesion and cell spreading (**Fig. 4G**, see **Supplementary Table S9, 10** for most downregulated and upregulated GO terms). Next, we focused on the ECM-related gene expression and FAK signaling. Having observed prominent shifts in terms related to cell-ECM interaction within the transcriptome and proteome of *NELL1* KO sarcoma cells, we next examined specific misexpressed ECM transcripts within *NELL1* KO cells (**Fig. 4H**). Analysis showed significant decrease in expression across several ECM components including for example *FN1* (Fibronectin 1), *LAMC2* (Laminin subunit gamma 2), and *COL6A3* (Collagen type VI alpha 3 chain) with *NELL1* gene deletion (**Fig. 4H**). Proteomic analysis confirmed a downregulation of FN1, as well as decreased phosphorylation of FAK and Src in *NELL1* KO cells (**Fig. 4I**). Thus, a transcriptomic-proteomic analysis suggested that deletion of *NELL1* in sarcoma cells alters the ECM profile, including prominent reductions in FN1 and Laminin, alterations in pathways associated with cell-ECM interaction, and a reduction FAK signaling activity.

### Matrix components can reverse phenotypic changes with NELL1 gene deletion

Having observed that *NELL1* gene deletion among 143B sarcoma cells resulted in deficits in key matricellular proteins and reduced FAK signaling activation, we next sought to confirm these findings across *in vivo* models. In a candidate fashion, fibronectin (FN1) and laminin (LAM) expression among 143B xenografts was assessed by immunofluorescent staining (**Fig. 5A**). Immunostaining showed diffuse FN1 and LAM immunoreactivity in control tumors, staining for both proteins was significantly reduced in *NELL1* KO tumors (**Fig. 5A**, 69.4% and 73.7% reduction in FN1 and LAM, respectively). Corresponding activation of FAK signaling was next accessed by immunofluorescent staining for FAK and pFAK across 143B sarcoma samples. Among *NELL1* KO sarcoma sections, decreased pFAK and retained FAK immunoreactivity was observed in comparison to control tumors (**Fig. 5B**, no significant differences in FAK, 58.3% reduction in pFAK, and 54.9% reduction in pFAK/FAK staining). These findings regarding altered sarcoma matrix were likewise observed in the murine spontaneous OS model induced by p53/Rb deletion (**Fig. 5C, D**). Here, sarcomas among *NELL1*^OC^ cKO animals demonstrated a 74.9% and 71.6% reduction in FN1 and LAM immunofluorescent staining across serial sections (**Fig. 5C**), associated with a 71.6% reduction in pFAK staining and 66.7% reduction in pFAK/FAK immunostaining in relation to control tumor tissues (**Fig. 5D**). This data further correlated *NELL1* gene deletion with shifts in sarcoma associated ECM composition and a corresponding reduction in FAK signaling.

**Figure 5.**
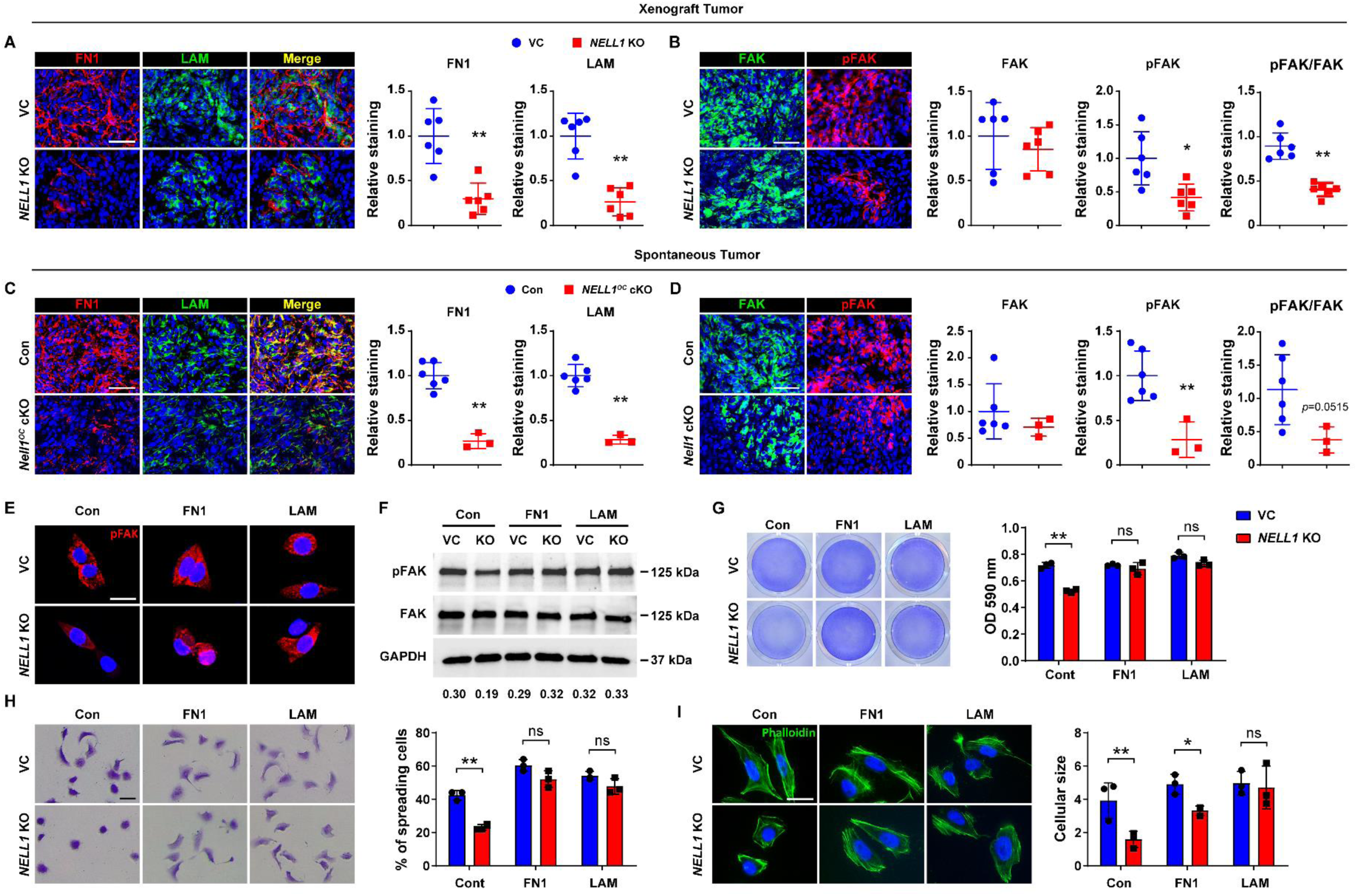
Sarcoma-associated ECM proteins positively regulate FAK signaling and adhesive features in *NELL1* KO sarcoma cells. **(A, B)** Immunohistochemical analysis of 143B OS cell implants among VC and *NELL1* KO tumor tissue, 28 d post-implantation. **(A)** Representative Fibronectin (FN1) and laminin (LAM) immunostaining (left) and quantification (right). **(B)** Representative FAK and pFAK immunostaining (left) and quantification (right). **(C, D)** Immunohistochemical analysis of mouse spontaneous OS tumor tissues from control (*p53*^fl/fl^;*Rb*^fl/fl^;OC-Cre) or *Nell1*^OC^ cKO (*p53*^fl/fl^;*Rb*^fl/fl^;*Nell1*^fl/fl^;OC-Cre) animals. **(C)** Representative FN1 and LAM immunostaining (left) and quantification (right). **(D)** Representative FAK and pFAK immunostaining (left) and quantification (right) among control and *Nell1*^OC^ cKO tumors. **(E-I)** Effects of sarcoma ECM components fibronectin (FN1) or laminin (LAM) on 143B OS cells with or without *NELL1* KO. Comparisons made to vector control (VC) treated cells. Pre-coated wells include FN1 (10 μg) and LAM (10 μg). **(E)** Representative pFAK immunostaining at 3 h post-seeding. **(F)** Western blot of pFAK and FAK at 5 h post-seeding. Ratio of pFAK/FAK listed below. **(G)** Attachment assay assessed by crystal violet staining (left) and quantification (right) at 5 h post-seeding. **(H)** Cell spreading assay assessed by crystal violet staining (left) and quantification (right) at 3 h. **(I)** Representative phalloidin staining (left) and quantification (right) at 3 h. Scale bar: 50 µm (**A-D**), and 20 µm (**E, H, I**). In (A-D), dots in scatterplots represent values from individual animal measurements, while mean and one SD are indicated by crosshairs and whiskers. In (E-G), data shown as mean ± 1 SD, with dots representing individual wells, performed in triplicate experimental replicates. \**P*<0.05; \*\**P*<0.01. ns: non-significant. A two-tailed student’s *t* test was used for all comparisons.

Next, the extent to which the matricellular proteins FN1 and LAM could restore FAK signaling and alter cellular behavior among *NELL1* KO 143B cells was assessed (**Fig. 5E-I**). Here, cell culture flasks were pre-coated with FN1 or LAM (5 μg/cm^2^). First, FAK activation was assessed in *NELL1* KO cells by pFAK immunocytochemistry and western blot (**Fig. 5E, F**). Results confirmed a decrease in FAK phosphorylation among *NELL1* KO cells in relation to vector control, which was observed to be restored by supplementation with FN1 or LAM coating (**Fig. 5E**). This finding was further validated by western blot and semi-quantitation (**Fig. 5F**, **Supplementary Fig. S8**). Further, restoration of FAK signaling activity with either FN1 or LAM coating led to alterations in cell attachment profiles among *NELL1* KO sarcoma cells. Here, complete restoration of attachment rate was observed among *NELL1* KO sarcoma cells on FN1 or LAM substrates (**Fig. 5G**). Correspondingly, a similar increase in adhesion-dependent cell spreading (**Fig. 5H**) and cytoskeletal size (**Fig. 5I**) was observed among *NELL1* KO sarcoma cells when placed on either matrix component. In sum, these data indicate that alterations in the sarcoma matrix composition mediate FAK signaling and tumor cell attachment potential, and implicate NELL1 as an important regulator of this process.

### NELL1 expression typifies an aggressive phenotype in human osteosarcoma

Finally, the critical role of NELL1 signaling in primary human osteosarcoma cells and tissues was examined. First, patient-derived human osteosarcoma cells (HuOS) were isolated and extensive karyotypic abnormalities were observed in the resulting cell isolates (**Fig. 6A, B**). Further, flow cytometry analysis confirmed absence of endothelial and hematopoietic markers (CD31, CD45) and presence of mesenchymal markers (CD44, CD73, CD90 and CD105) and among cell isolates (**Fig. 6C**, **Supplementary Fig. S9**). In addition, all isolated osteosarcoma cells underwent both osteogenic and chondrogenic differentiation (**Fig. 6D**). Basal expression of *NELL1* was observed across all HuOS cells (**Fig. 6E**). Gene deletion studies were then performed among patient-derived OS cells using CRISPR/Cas9. Knockout was confirmed using qRT-PCR (**Fig. 6F**, 88.1% reduction). In comparison to vector control, multiple cellular effects were observed among HuOS, including reduced proliferation (**Fig. 6G**. 46.3% reduction at 72 h), attachment (**Fig. 6H**. 34.9% and 24.7% reduction at 3 and 5 h, respectively), invasion (**Fig. 6I**, 47.2% reduction at 24 h), and tumorsphere formation (**Fig. 6J**, 69.7% reduction at 5 d).

**Figure 6.**
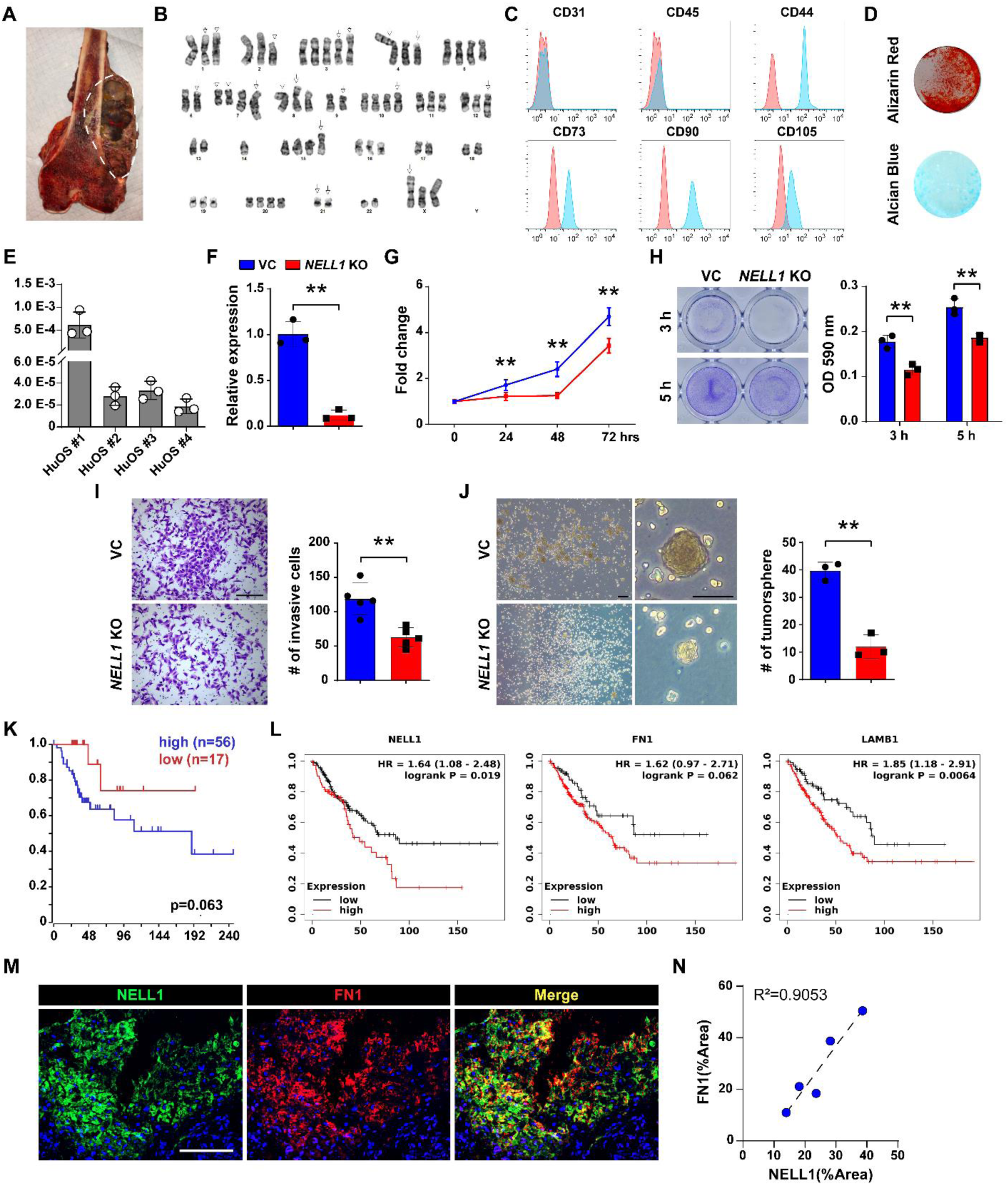
*NELL1* regulates invasive potential in primary human OS cells and is associated with disease progression in human patients. (**A**) Representative image of bisected osteosarcoma sample. White dashed lines indicate tumor. (**B**) Representative karyotype of primary osteosarcoma cells. (**C**) Representative flow cytometry analysis of patient-derived primary osteosarcoma cells. Frequency of expression is shown (blue) in relation to isotype control (red). (**D**) Representative Alizarin red (AR) at d 10 of osteogenic differentiation, and Alcian blue staining at d 14 of chondrogenic differentiation among human OS cells. (**E**) *NELL1* expression in primary osteosarcoma cells by qRT-PCR. (**F-J**) Effects of *NELL1* gene deletion in primary osteosarcoma cells. (**F**) Confirmation of *NELL1* gene deletion efficiency by qRT-PCR. MTS proliferation assay among *NELL1* KO primary osteosarcoma cells at 24, 48, and 72 h. (G) Attachment assay assessed by crystal violet staining and quantification, at 3 and 5 h. (**I**) Transwell invasion assay and quantification, at 24 h. Representative images with number of invasive cells shown. Scale bars: 500 µm. (**J**) Tumorsphere formation assay and quantification at 5 d. (**K**) Prognostic value of *NELL1* in patients with high grade osteosarcoma. Kaplan–Meier curve generated by R2 genomics analysis and visualization platform. (**L**) Prognostic value of *NELL1*, *FN1*, and *LAMB1* in high grade sarcoma patients. Kaplan–Meier curve generated by Kaplan-Meier (KM) plotter pan-caner RNA seq. (**M**) Representative NELL1 and Fibronectin immunofluorescent staining in biopsy samples of high grade conventional osteosarcoma. (**N**) Correlation of NELL1 and Fibronectin staining across N=5 biopsy samples of high grade conventional osteosarcoma. Scale bars: 100 µm. Data shown as mean ± 1 SD, and represent triplicate experimental replicates. \**P*<0.05; \*\**P*<0.01. A two-tail Student’s t test was used for two group comparisons.

Two existing datasets were used to first examine whether high expression of NELL1 serves as a prognostic marker in OS (**Fig. 6K, L**). Among a mixed population of 73 cases of high grade osteosarcoma, high *NELL1* transcripts were associated with a reduced overall survival (**Fig. 6K**, median overall survival of 34 months in NELL1^high^ versus 60 months in NELL1^low^ cohort).

Analyses performed in 259 cases of high grade sarcomas of soft tissue and bone showed similarly that transcripts of *NELL1, FN1,* or *LAMB1* were all associated with decreased overall survival (**Fig. 6L**, median overall survival of 49, 23, and 54 months in NELL1^high^ versus 86, 38, and 90 months in NELL1^low^ cohort, respectively). Finally, 5 biopsies of conventional osteosarcoma prior to adjuvant chemotherapy were assessed for the expression of NELL1 and fibronectin (**Fig. 6M, N, Supplementary Table S11**). Results demonstrated immunoreactivity for both NELL1 and FN1 across all samples, a high degree of co-localization between antigens within biopsy tissue, and a relative correlation between the degree of NELL1 and FN1 immunostaining across samples ((**Fig. 6M, N**). Collectively, *NELL1* is an OS related protein essential for sarcoma cell function via regulation of cell-matrix interactions, and represents a poor prognostic indicator across human sarcomas.

## Discussion

In this study, we demonstrated that the sarcoma associated protein NELL1 plays an integral role in positively regulating both sarcomagenesis as well as features of aggressive sarcoma behavior on cellular and tumoral levels. In part, this is explained by a modulation of the sarcomatous ECM, which in turn modulates cell-ECM interaction and FAK/Src signaling activation.

The sarcomatous ECM displays distinct histologic, proteomic, and biomechanical properties ^42–45^. In some instances, sarcomatous ECM proteins have been linked to metastasis ^44^, subsequent growth of metastatic foci ^43^, or chemo-resistance ^15, 19, 42^. Bone and soft tissue sarcomas are exceptionally diverse ^46, 47^, and even within osteosarcoma the matrix components may vary widely in composition ^15, 16^. Nevertheless, NELL1 expression appears to be present across the histologic spectrum of human osteosarcoma ^31^. Prior observations have found NELL1 to be a key protein in the regulation of matrix components during morphogenesis, regeneration and tissue engineering. For example, mice globally deficient in *Nell1* have altered ECM protein expression with associated with skeletal developmental anomalies ^30, 48^. In the context of tissue engineering and regeneration, Qi *et al* showed recombinant NELL1 treatment induces expression of the structural matrix protein Type II Collagen and promotes the differentiation of chondrocytes ^48^. In other contexts, NELL1 enhances the production of the molecule dentin matrix protein 1 (DMP1) accompanied by induction of odontoblastic differentiation ^49^. Our study observed similarly striking regulation of sarcoma associated matrix proteins by NELL1, which among other findings altered FAK signaling activity and modulated sarcoma cell functional characteristics.

An important consideration toward the positive regulatory roles of NELL1 signaling in sarcoma disease progression is the promotion of angiogenesis. In tissue engineering contexts, NELL1 is known to promote angiogenesis during non-neoplastic bone formation through induction of VEGFA expression ^32, 33^. Here, *NELL1* gene deletion led to a down-regulation of VEGF signaling in sarcoma cells, and potent reduction in sarcoma-associated blood vessels. Indeed, VEGF expression correlates with prognosis in patients with sarcoma ^50^. Beyond changes in angiogenic factors, it is likely that altered ECM associated with *NELL1* KO may also have predisposed to lack of vascular ingrowth. NELL1 deficient sarcoma cells displayed changes in genes and proteins associated with ECM remodeling, including a significant downregulation in *MMP9.* Matrix metalloproteases (MMPs) are defined by their role in matrix degradation but also in their secondary stimulation of tumor vasculature ^51^. For instance, MMP9 renders normal islets into angiogenic islets by mobilizing VEGF during pancreatic carcinogenesis ^52^. Thus, changes in sarcoma-associated angiogenesis could be a result of combinatorial factors including NELL1 signaling induced transcription of angiogenic factors as well as altered ECM remodeling.

NELL1 is by no means solely expressed by osteosarcoma. Here, we observed NELL1 expression to serve as a prognostic biomarker among high grade sarcomas of bone (osteosarcoma and Ewings sarcoma), or even mixed diagnostic population of high grade sarcomas of bone and soft tissue. This is consistent with a prior study which implicated *NELL1* in migration and invasion of rhabdomyosarcoma (RMS) cells, and negatively correlated *NELL1* expression with RMS prognosis ^53^. Interestingly, a clear divide seems to be present in the literature between the potential roles of NELL1 in epithelial versus mesenchymal malignant tumors. For example, a GWAS study of non-small cell lung cancer (NSCLC) identified NELL1 as a tumor suppressor gene ^54^. The protective role of NELL1 was confirmed by functional studies conducted in colon cancer, renal cell carcinoma, lung cancer and esophageal adenocarcinoma cell lines and models ^55–58^. Among the numerous differences between carcinomas and sarcomas, our study suggests that the matrix itself may be an important distinguishing factor.

Sarcoma cells are often loosely and often haphazardly arranged their sarcomatous ECM ^59^, while epithelial cancer cells are closely interacting cells often adherent to one another, but by and large without mesenchymal matrix production. These key cytologic and histologic features serve as future catalysts toward the understanding of the matrix regulating protein NELL1 as a differential disease modifier in carcinoma and sarcoma.

In combination, our aggregate data suggest that NELL1 is a novel protein highly expressed among human osteosarcoma which positively regulates multiple aspects of OS disease progression. Use of gene editing technology, or alternatively small molecule-mediated disruption of NELL1 signaling, may represent an adjunctive tool to improve survival in osteosarcoma and other sarcoma subtypes.

## Materials and Methods

### Mice

All animal experiments were performed according to approved protocols (MO21M112) of the Animal Care and Use Committee (ACUC) at Johns Hopkins University (JHU). NOD scid mice (Stock No: 001303) were purchased from The Jackson Laboratory (Bar Harbor, ME, USA). OC-Cre animals were a kind gift from the Clemens laboratory at JHU (JAX Stock No. 019509). *p53*^fl/fl^;*Rb*^fl/fl^ mice were a kind gift from the Jones laboratory at University of Utah (JAX Stock No. 008462 and 026563). *Nell1*^fl/fl^ mice were supplied by the Ting/Zhang laboratories at UCLA ^48^. In order to achieve spontaneous osteosarcoma formation, *p53*^fl/fl^;*Rb*^fl/fl^ (control) mice or *p53*^fl/fl^;*Rb*^fl/fl^;*Nell1*^fl/fl^ (*Nell1*^OC^ cKO) mice were crossbred with OC-Cre animals. Rates of OS tumor formation within *p53*^fl/fl^;*Rb*^fl/fl^;OC-Cre animals were in line with prior reports ^37^. Analyses were performed by investigators blinded to the mouse genotype.

### Cell isolation, culture and characterization

Human osteosarcoma cell lines were purchased from American Type Culture Collection (Manassas, VA), including 143B (ATCC®-CRL-8303™), Saos-2 (ATCC^®^ HTB-85^™^), HOS (ATCC^®^ CRL-1543^™^), KHOS/NP (ATCC® CRL-1544™), KHOS-312H (ATCC^®^ CRL-1546^™^), and G-292 (ATCC® CRL-1423™). Primary human OS cells were derived from human OS resection samples (n=4) under IRB approval at JHU with a waiver of informed consent. The pathologic diagnosis of high grade conventional osteosarcoma was confirmed by two independent bone pathologists (E.F.M. and A.W.J.). Immediately following resection, specimens of human OS were collected under sterile conditions, washed in PBS and dissected into small fragments (<1 mm^3^). Primary OS tumor cells were derived as per previous descriptions ^60^, and culture expanded for 3 to 5 passages before use. Characterization of primary tumor cells was performed by flow cytometry, *in vitro* differentiation and cytogenetic analyses. Karyotyping was performed by the Division of Molecular Pathology at JHU clinical laboratories. Human primary OS cells at passage 3-5 were used for characterization by flow cytometry ^61^, including mesenchymal (CD44, CD73, CD90, CD105), endothelial (CD31) and hematopoietic markers (CD45) (see **Supplementary Table S12** for all antibody information). Multilineage differentiation of tumor cells was performed for osteogenic and chondrogenic differentiation ^62^, with assessments of mineralization at 10 d and glycosaminoglycan at 14 d ^63–65^. Mouse bone marrow mesenchymal stromal cells (BMSCs) were isolated from tibial and femoral bone marrow of either *p53*^fl/fl^;*Rb*^fl/fl^ (DKO) or *p53*^fl/fl^;*Rb*^fl/fl^;*Nell1*^fl/fl^ (TKO) mice (n=4) under sterile conditions ^66, 67^. Briefly, bones were dissected from 8-12 week old female mice. Bone marrow was flushed, resuspended in PBS, and filtered through a 70-mm filter. Bone marrow cells were cultured in 10-cm culture dishes and incubated at 37 °C with 5% CO2 in a humidified chamber. After 3 h, non-adherent cells were removed. All cells were cultured in Dulbecco’s Modified Eagle Medium (DMEM, Gibco, Grand Island, NY) supplemented with 15% fetal bovine serum (FBS, Gibco), 100 U/ml penicillin and 100 µg/ml streptomycin (Gibco) in a humidified incubator with 5% CO_2_ at 37°C.

### NELL1 gene deletion and isolation of cell clones

CRISPR/Cas9 gene deletion of *NELL1* in human OS cell lines and primary cells was achieved using a plasmid with a guide RNA (gRNA) sequence combined with U6gRNA-Cas9-2A-RFP vector (Sigma-Aldrich). The specific gRNA sequence of knockout and negative control plasmids were as in **Supplementary Table S13**. The transient plasmid DNA transfection was performed using TransIT®-LT1 Transfection Reagent (Mirus, Madisom, WI). Briefly, cells were plated at an initial density of 3×10^5^ cells/ml in 6-well cell culture plates and incubated for 24 h. Cells were then transfected with TransIT-LT1 Reagent-plasmid DNA complex containing 250 μl of Opti-MEM Reduced-Serum Medium (Gibco), 2.5 μg plasmid DNA and 7.5 μl TransIT-LT1 Reagent and incubated for 72 h. Next, *NELL1* KO single cell colonies were established by FACS. Viable RFP-positive cells from *NELL1* KO-transfected cells and control-transfected cells were sorted into wells of a 96-well microtiter plate by using a Dako Cytomation MoFlo cell-sorter (Beckman, Indianapolis, IN). For single cell deposition, cells were sorted using the 70-mm nozzle with the sheath pressure set at 70 PSI using the sort precision mode set at single cell.

After confirmation of gene deletion by qRT-PCR, n=6 colonies were expanded for use. Finally, to achieve *Nell1* gene deletion of in murine cells, BMSCs derived from either *p53*^fl/fl^;*Rb*^fl/fl^ (DKO) or *p53*^fl/fl^;*Rb*^fl/fl^;*Nell1*^fl/fl^ (TKO) mice and were treated with either Ad-GFP-2A-iCre (Vector Biolabs, Cat No. 1772) or Ad-CMV-GFP (Vector Biolabs, Cat No. 1060) at 200 MOI.

### T7 Endonuclease I Assay

*NELL1* KO and control-transfected cells were collected for the detection of the mutation using the T7 Endonuclease I Assay Kit (Genocopoeis, Rockville, MD). DNA was extracted from the samples using Quick-DNA™ Miniprep Kit (Zymo Research, Irvine, CA). See **Supplementary Table S13** for primer information. The PCR reactions were performed according to the manufacturer’s protocol. The PCR product was then digested using T7E1 enzyme at 37°C for 60 min, followed by separation using 1.5% agarose gel electrophoresis.

### In vitro functional assays

Firstly, cell proliferation was assessed using the CellTiter 96® AQueous Non-Radioactive MTS Cell Proliferation Assay (Promega, Madison, WI) ^62, 68^. 2 × 10^3^ cells were plated in 96-well plates and incubated for 24, 48, and 72 h. The optical density at 490 nm was measured on a microplate spectrophotometer (BioTek, Winooski, VT). Second, cell attachment was measured by crystal violet staining ^69^. 2 × 10^5^ cells were plated in 24-well plates and incubated for 1, 3 and 5 h. Cells were washed with cold PBS and fixed with 100% methanol, then stained with 0.5% crystal violet solution for 10 min at RT. Stained cells were lysed in 10% acetic acid and absorbance was measured at 570 nm. In select experiments, *Nell1* signaling was rescued by adenoviral NELL1 (Ad-*NELL1*) or control (Ad-LacZ). In select experiments, tissue culture plates were pre-coated with 5 µg/cm^2^ of either Fibronectin (Sigma-Millipore) or Laminin (Sigma-Millipore). Thirdly, migration assays were carried out using the Culture-Insert 2 Well (Ibidi, Martinsried, Germany) ^63^. 4 × 10^4^ cells were added into each well and incubated overnight. Next, the insert wells were gently removed and the plate replenished with growth medium. The cells were incubated for 12 and 24 h and photographs were taken under an inverted microscope (Olympus, Tokyo, Japan). The width of the cell-free gap was measured by using ImageJ software (Version 1.49 v, NIH, Bethesda, MD).

Fourth, invasion assays were carried out using Corning® BioCoat™ Matrigel® Invasion Chambers (Corning, Bedford, MA). Briefly, 2.5 × 10^5^ cells in serum-free medium were added to the upper chamber, while 500 μl growth medium was added to the lower chamber. After 24 h incubation, cells were fixed with cold 100% methanol and stained with 0.5% crystal violet solution. Photographs of five random regions were taken under an inverted microscope (Olympus) and the number of invaded cells was enumerated. Fifth, tumorsphere assays were performed using primary OS cells with or without *NELL1* KO which were trypsinized to generate a single cell suspension and seeded into 6-well ultralow attachment plates (Corning) at a density of 5 × 10^5^ cells, followed by culturing in defined sphere medium ^70, 71^. Spheres were imaged on day 7 using an inverted microscope (Olympus) and the number of spheres were manually counted.

### qRT-PCR

Total RNA was extracted from cultured cells of equal passage number and density using TRIzol Reagent (Invitrogen, Carlsbad, CA) according to the manufacturer’s instructions. 1 μg of total RNA was used for reverse transcription with iScript cDNA synthesis kit (Bio-Rad). Real-time PCR was performed using SYBR Green PCR Master Mix (Thermo Scientific, Waltham, MA). Relative gene expression was calculated using a 2^-ΔΔCt^ method by normalization with GAPDH. The primer sequences were listed in **Supplemental Table S13**.

### Western blot

Cells were lysed in RIPA buffer (Thermo Scientific) with protease inhibitor cocktail (Cell Signaling Technology, Danvers, MA, USA). Proteins concentration were determined by BCA assay (Thermo Scientific). Cell lysates were separated by SDS-polyacrylamide gel electrophoresis and transferred onto a nitrocellulose membrane. Protein extraction was blocked with 5% bovine serum albumin and incubated with primary antibodies at 4°C overnight. Finally, membranes were incubated with a horseradish-peroxidase (HRP)-conjugated secondary antibody and detected with ChemiDoc XRS+ System (Bio-rad). The antibodies were listed in **Supplemental Table S12**.

### Total RNA Sequencing

Gene expression was detected by total RNA sequencing using the Illumina NextSeq 500 platform (Illumina, San Diego, CA). Briefly, total RNA was extract from clonal 143B cells with or without *NELL1* KO by Trizol (Life Technologies Corporation, Gaithersburg, MD). The total RNA samples were sent to the JHMI Deep Sequencing and Microarray core (JHU) for sequencing. Data analysis were performed using software packages including CLC Genomics Server and Workbench (RRID:SCR_017396 and RRID:SCR_011853), Partek Genomics Suite (RRID:SCR_011860), Spotfire DecisopnSite with Functional Genomics (RRID:SCR_008858), and QIAGEN Ingenuity Pathway Analysis (RRID:SCR_008653).

### Reverse phase protein analysis

Protein lysates were isolated from control and *NELL1* KO clonal 143B cells (4 biological replicates). Modified RIPA buffer containing proteinase and phosphatase inhibitor was used for the lysis. Cellular proteins were denatured by 1% SDS with β-mercaptoethanol and analyzed by the Reverse Phase Protein Array (RPPA) Core at MD Anderson Cancer Center (https://www.mdanderson.org/research/research-resources/core-facilities/functional-proteomics-rppa-core.html). Briefly, protein lysates were diluted in five two-fold serial dilutions (undiluted, 1:2, 1:4, 1:8; 1:16) and arrayed on nitrocellulose-coated slides (Grace Bio Lab). Each slide was probed with a validated primary antibody plus a biotin-conjugated secondary antibody. The RPPA slides for 453 unique antibodies were scanned, analyzed, and quantified using a customized-software to generate spot intensity. Relative protein levels for each sample were determined by interpolating each dilution curve produced from the densities of the 5-dilution sample spots using a standard curve for each antibody. Relative protein levels are designated as log2 values. The protein concentrations of each set of slides were then normalized for protein loading ^72, 73^.

### Osteosarcoma implantation and assessments

All animal studies were performed with institutional ACUC approval within Johns Hopkins University, complying with all relevant ethical regulations. Experimental details and animal numbers are also summarized in **Supplementary Table S14**. For all 143B OS cell implantation, 8-10 week old, male and female, NOD scid mice were used. 1 × 10^6^ clonal 143B cells with or without *NELL1* KO in 50 μl PBS were injected subperiosteally within the proximal tibia metaphysis. Tumor size was measured by caliper twice weekly for 4 wks, and tumor volume was calculated ^74^. Experiments were replicated with a second cell clone. Magnetic resonance imaging (MRI) was performed as previously described ^75^ at 14 and 28 d post injection, and tumor size was calculated using ImageJ. Primary tumors and lungs were harvested at 28 d post-injection for histologic analysis. In a separate study, overall survival was evaluated from the first day of tumor cell injection until death using the Kaplan-Meier estimate. In a separate study, 143B cells with or without *NELL*1 knockout were injected systemically by tail vein (1.5 × 10^6^ clonal 143B cells in 100 μl PBS). Mice were monitored three time weekly and sacrificed 28 d after injection for assessment of pulmonary burden of disease. In a separate study, 1 × 10^6^ murine BMSCs in 50 μl PBS (*p53*^fl/fl^;*Rb*^fl/fl^ or *p53*^fl/fl^;*Rb*^fl/fl^;*Nell1*^fl/fl^ genotypes 2 d after Ad-Cre mediated transformation) were injected subperiosteally within the proximal tibia metaphysis.

Tumor formation was monitored three times weekly and tumor free survival was recorded. Samples were harvested 28 d after tumor formation was observed. Analyses were performed by investigators blinded to the tumor type.

### Spontaneous osteosarcomagenesis and assessments

The spontaneous osteosarcoma mice model is based on osteoblast-restricted deletion of p53 and pRb ^35, 36^. Either *p53*^fl/fl^;*Rb*^fl/fl^ mice or *p53*^fl/fl^;*Rb*^fl/fl^;*Nell1*^fl/fl^ mice were crossed with *p53*^fl/+^;*Rb*^fl/+^;OC-Cre mice or *p53*^fl/+^;*Rb*^fl/+^;*Nell1*^fl/+^;OC-Cre mice, respectively. The offspring with genotype of *p53*^fl/fl^;*Rb*^fl/fl^; OC-Cre (control) and *p53*^fl/fl^;*Rb*^fl/fl^;*Nell1*^fl/fl^;OC-Cre (*Nell1*^OC^ cKO) were used in this study. Tumor formation was monitored three times weekly and tumor free survival was recorded. Tumors were harvested 28 d after tumor formation was observed. Analyses were performed by investigators blinded to the mouse genotype.

### Histology and Immunohistochemistry

Primary tumor samples and lungs were fixed in 4% paraformaldehyde (PFA) at 4°C for 24 h and decalcified in 14% EDTA (Sigma-Aldrich) for up to 56 d at 4°C. Samples were then cryoprotected in 30% sucrose overnight at 4°C and embedded in optimal cutting temperature compound (OCT, Tissue-Tek 4583, Torrance, CA) for cryosectioning at 10 µm thickness. Routine H&E staining was performed on the entire lung tissues sectioned in a coronal plane to assess pulmonary burden, and number of metastasis foci were manually counted. For immunohistochemistry, sagittal sections of primary or metastatic tumors were washed in PBS × 3 for 10 min and permeabilized with 0.5% Triton-X for 30 min. Next, 5% normal goat serum (S-1000, Vector Laboratories, Burlingame, CA) was applied for 30 min, then incubated in primary antibodies overnight at 4°C. The following day, slides were washed in PBS, incubated in the secondary antibody for 1 h at 25°C, and then mounted with DAPI mounting solution (Vectashield H-1500, Vector Laboratories). Digital images of these sections were captured with 10-100 × objectives using upright fluorescent microscopy (Leica DM6, Leica Microsystems Inc., Buffalo Grove, IL). Analyses were performed by investigators blinded to the sample identification.

### Statistical analysis

Results are expressed as the mean ± 1 SD. A Shapiro-Wilk test for normality was performed on all datasets. Homogeneity was confirmed by a comparison of variances test. Parametric data was analyzed using either a Student’s t test for a two group comparison, or one-way analysis of variance when more than two groups were compared, followed by a post hoc Tukey test (Graphpad Software 7.0). \**P*<0.05 and \*\**P*<0.01 were considered significant. For *in vivo* 143B implantation studies, the sample size was calculated based on an anticipated effect size of 2.0 based on our *in vitro* studies comparing *NELL1* KO and vector control. For this scenario, with eight replicates per group will provide 80% power to detect effect sizes of at least 1.5, assuming a two-sided 0.05 level of significance.

## Acknowledgments

Support to AWJ is provided by American Cancer Society (Research Scholar Grant, RSG-18-027-01-CSM), NIH/NIAMS (R01 AR070773, R01 AR079171), Department of Defense (USAMRAA W81XWH-18-1-0336, W81XWH-18-1-0121, W81XWH-20-1-0795, W81XWH-20-1-0302), and the Maryland Stem Cell Research Foundation. The content is solely the responsibility of the authors and does not necessarily represent the official views of the National Institute of Health or Department of Defense. We thank the JHU microscopy core facility, JHMI deep sequencing and microarray core facility, division of molecular pathology and Hao Zhang within the JHU Bloomberg Flow Cytometry and Immunology Core.

## Author contributions

Conception and design: Q.Q., A.W.J.; funding and final manuscript approval: A.W.J.; acquisition, analysis, and interpretation of data: Q.Q., M.G.S., R.J.T., L.C., K.T., and X.Z.; donation of clinical samples: C.D.M., and E.F.M.; manuscript preparation: Q.Q., and A.W.J.

## Supplementary Materials

**Fig. S1.**
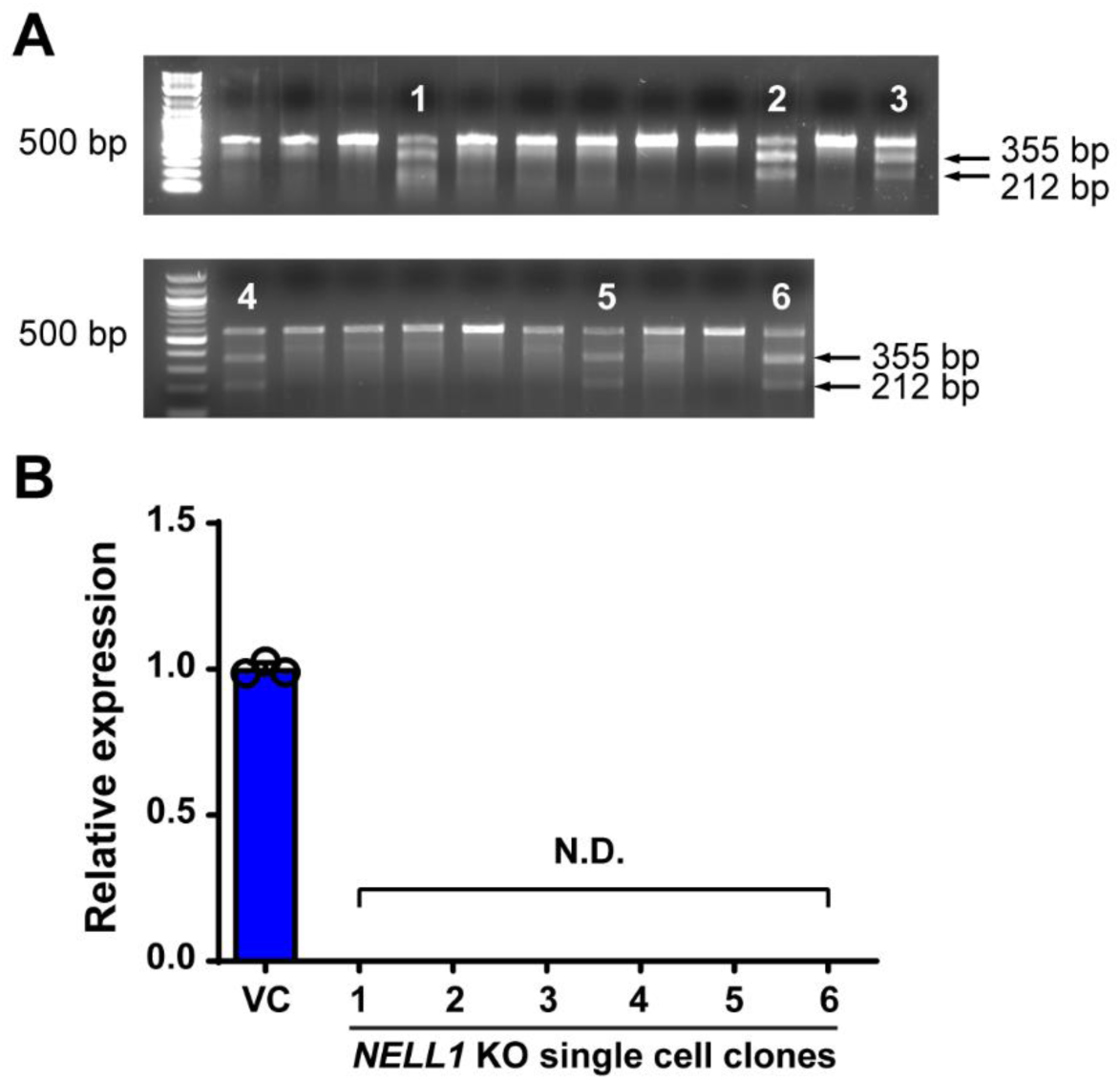
Validation of CRISPR-Cas9 mediated *NELL1* KO efficiency. Confirmation of *NELL1* knockout (KO) in six single cell clones by (**A**) T7 endonuclease I assay and (**B**) qRT-PCR. (**A**) The arrows indicate the cleaved PCR products corresponding to 355 and 212 bp respectively. (**B**) Relative expression of *NELL1* in six clones compared with vector control (VC). Data shown as mean ± 1 SD, with dots representing individual datapoints. All experiments performed in triplicate replicates, with results from a single replicate shown. ND: Not detected.

**Fig. S2.**
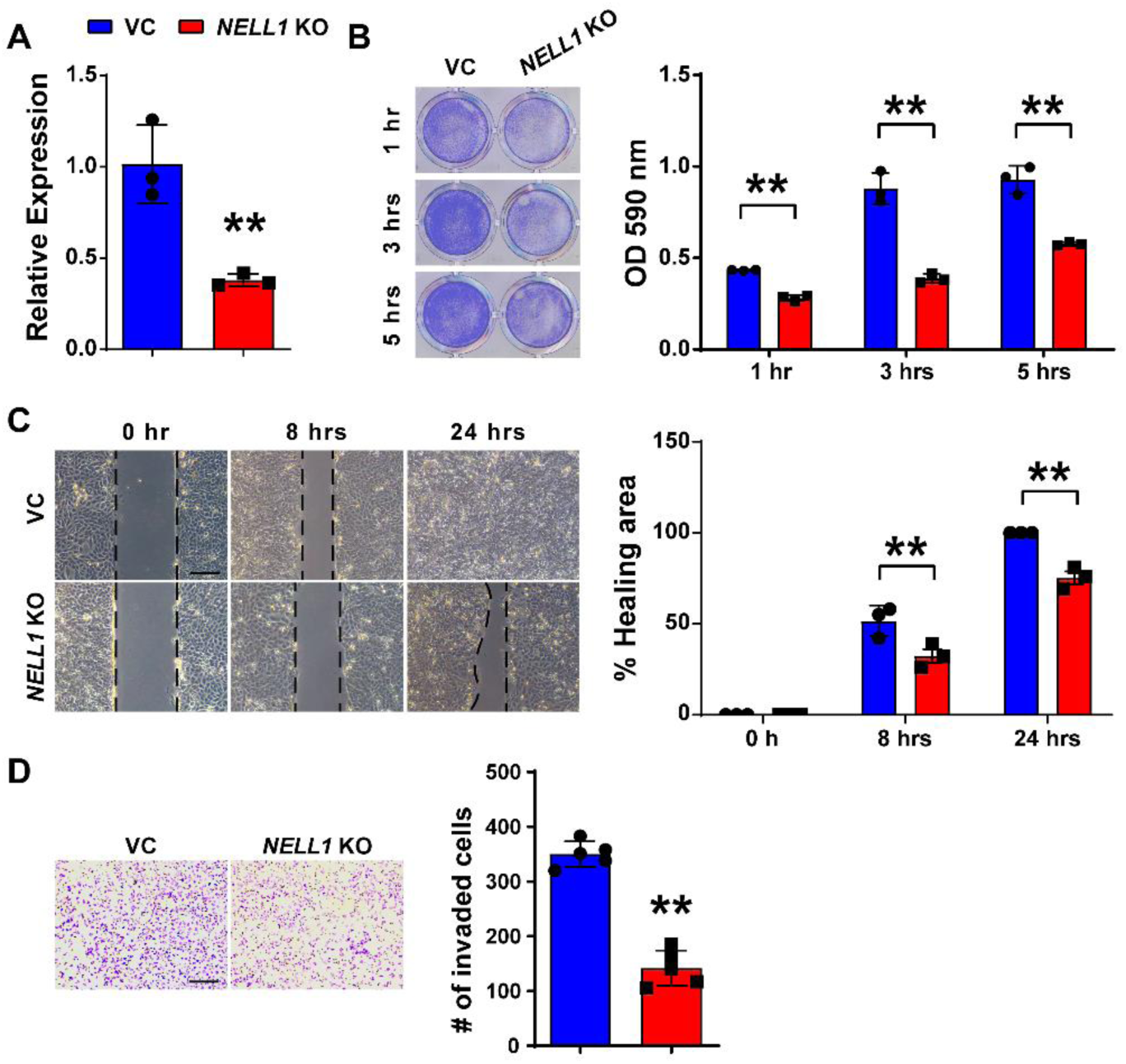
CRISPR-Cas9 mediated *NELL1* KO in polyclonal 143B cells. (**A**) Confirmation of *NEL*L1 gene deletion efficiency by qRT-PCR, in comparison to VC. (**B-D**) Effects of CRISPR-mediated *NELL1* gene deletion in polyclonal 143B OS cells. (**B**) Attachment assay as assessed by crystal violet staining (left) and quantification (right) with or without *NELL*1 KO (1-5 h). (**C**) Scratch wound healing assay with or without *NELL1* KO. Representative images (left) with percentage wound healing (right) (8 and 24 h). **(D)** Transwell invasion assay with or without *NELL1* KO (24 h). Representative images (left) with number of invaded cells (right). Data shown as mean ± 1 SD, with dots representing individual datapoints. All experiments performed in triplicate replicates, with results from a single replicate shown. \*\**P*<0.01. Scale bars: 500 µm. A two-tail Student’s *t* test was used for two group comparisons.

**Fig. S3.**
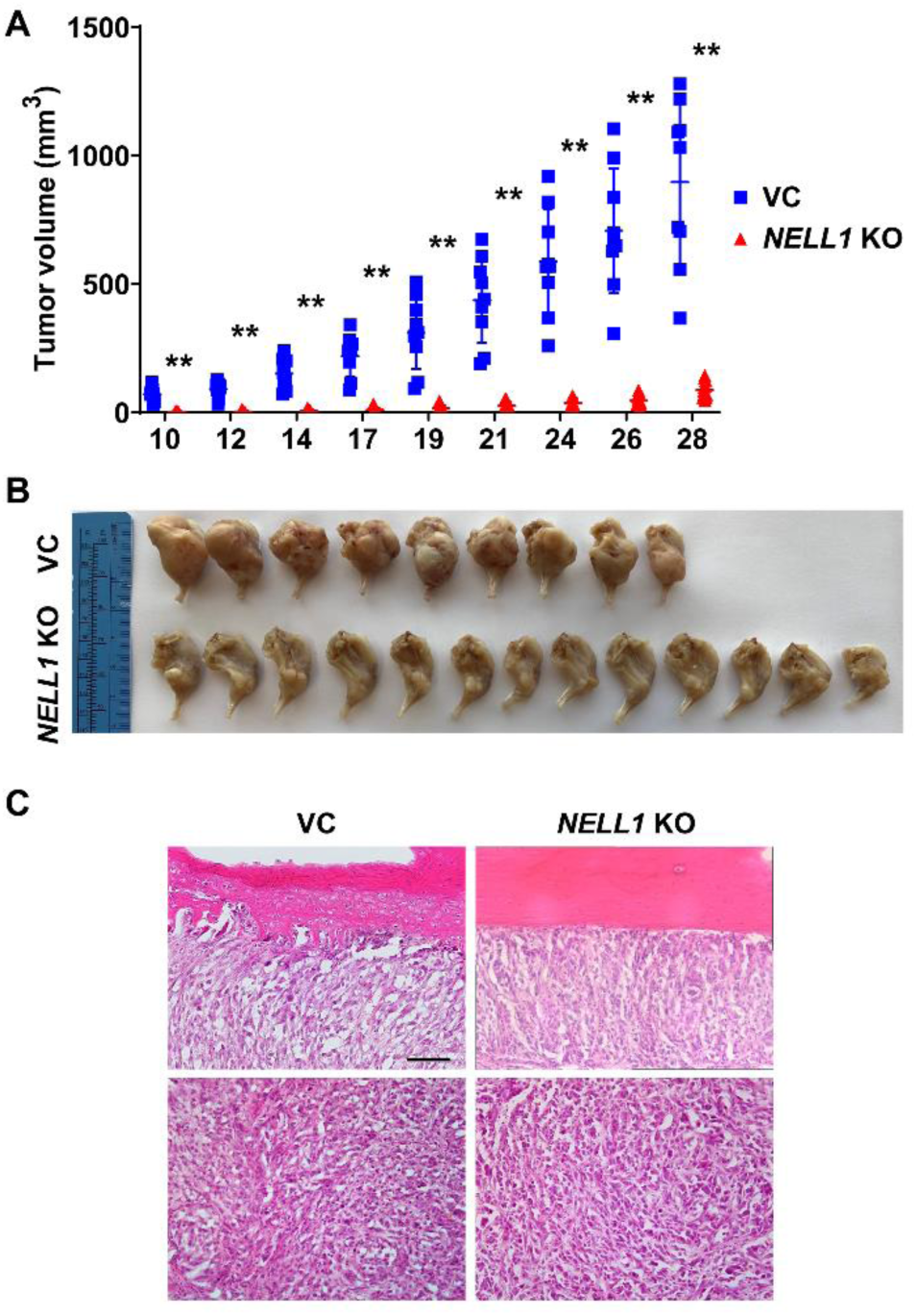
*NELL1* gene deletion mitigates OS disease progression. Biologic replicate of data shown in **Fig. 3B**. Orthotopic implantation of *NELL1* KO or vector control (VC) 143B cells within the proximal tibial metaphysis in NOD-Scid mice (n=9 VC implants and n=13 *NELL1* KO implants, 1 × 10^6^ cells per implant). **(A)** Tumor volume, calculated by caliper measurements until 28 d post-injection. **(B)** Gross pathology of all tumors. **(C)** Representative histologic appearance by routine H&E staining, adjacent to the tibial cortex (above) and a representative central area of tumor (below). Scale bars: 200 µm. Dots in scatterplots represent values from individual implants, while mean and one SD are indicated by crosshairs and whiskers. \*\**P*<0.01. A two-tailed student’s t test was used for all comparisons.

**Fig. S4.**
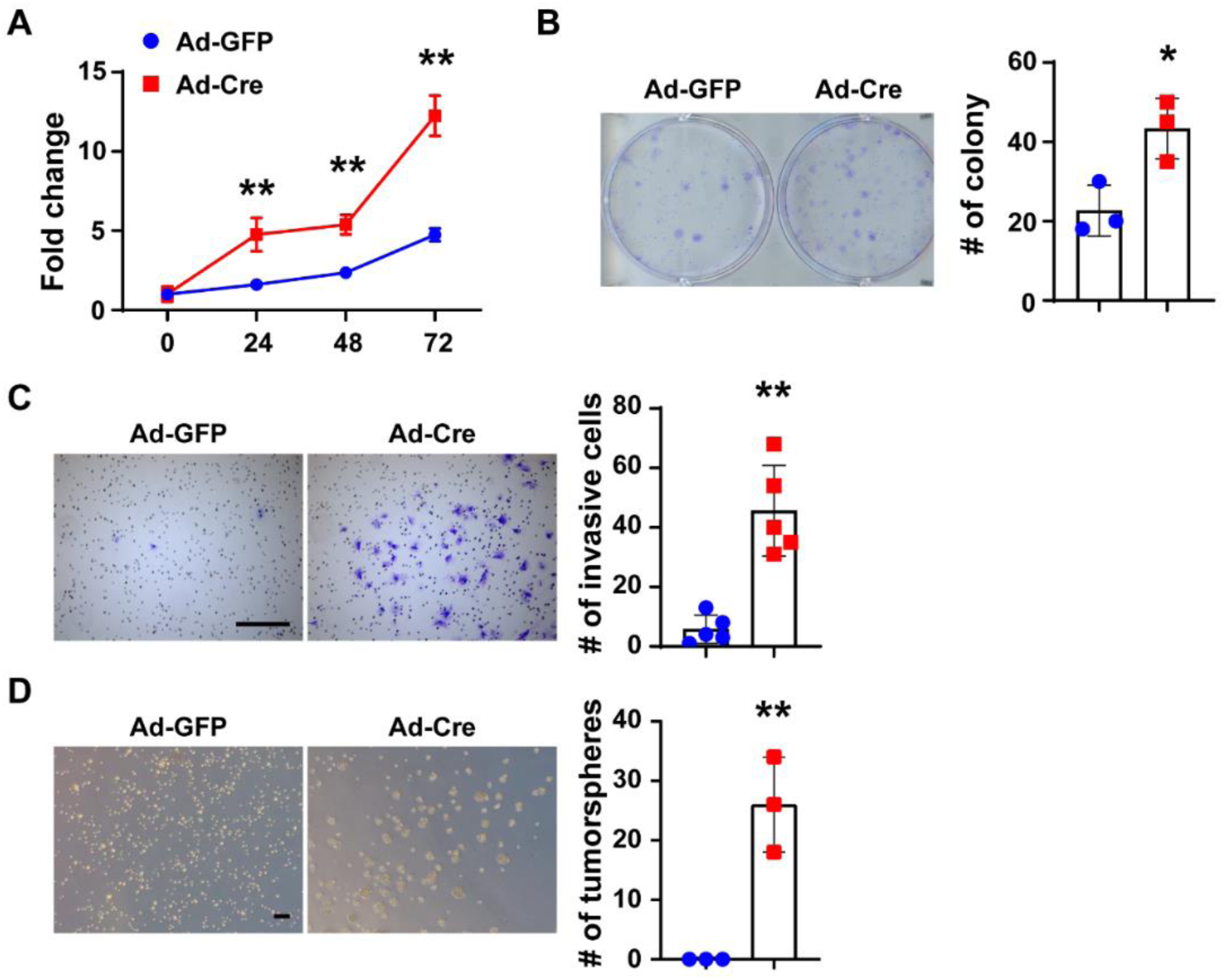
Cellular behavior after adenovirus-mediated p53/Rb gene deletion in mouse BMSCs. BMSCs were isolated from *p53*^fl/fl^;*Rb*^fl/fl^ mice and transfected with either Ad-GFP or Ad-Cre (200 MOI). **(A)** MTS proliferation assay at up to 72 h. **(B)** Colony-forming unit (CFU) assay assessed by crystal violet staining (left) and quantification (right) at 10 d. **(C)** Invasion assessed by transwell assay, crystal violet staining (left) and quantification (right) at 24 h. **(D)** Tumorsphere formation assay (left) and quantification (right) at 5 d. Scale bars: 100 µm. Data shown as mean ± 1 SD, and represent triplicate experimental replicates. Dots in scatterplots represent values from individual wells. \**P*<0.05; \*\**P*<0.01. A two-tailed student’s t test was used for all comparisons.

**Fig. S5.**
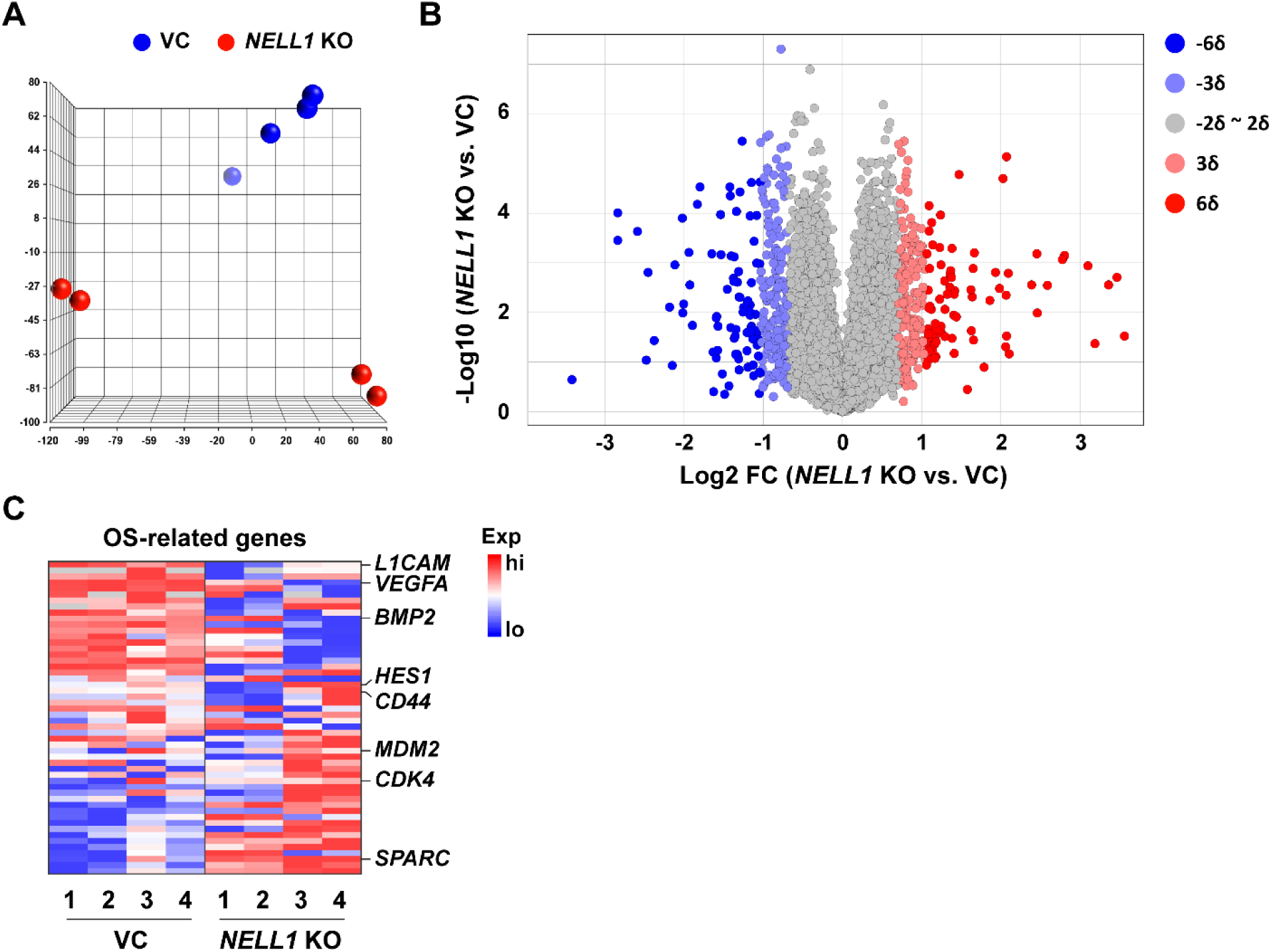
Additional transcriptomic analysis of clonal 143B cells with or without *NELL1* gene deletion. **(A)** Principal component analysis among VC and *NELL1* KO osteosarcoma cells. **(B)** Volcano plot of 54,174 protein coding genes expressed among VC and *NELL1* KO 143B cells. Red dots indicate >2SD increase among *NELL1* KO cells. Blue dots indicate >2SD decrease among *NELL1* KO cells. **(C)** Heat map indicating OS-related gene expression among VC and *NELL1* KO osteosarcoma cells, with some representative genes listed. See **Supplementary Table S8** for a detailed gene list. n=4 biological replicates of VC and *NELL1* KO 143B OS cells.

**Fig. S6.**
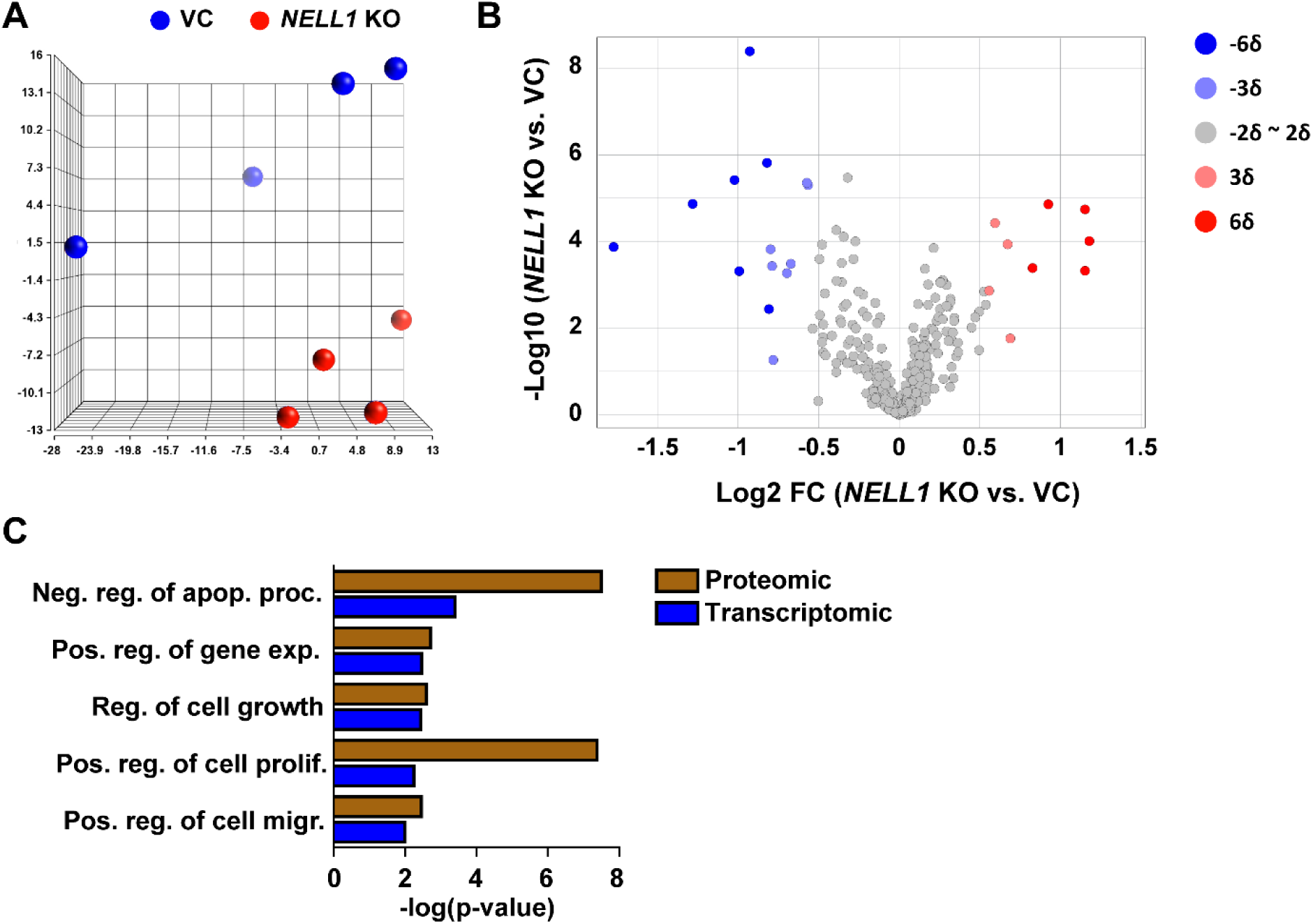
Additional proteomic analysis of clonal 143B cells with or without *NELL1* gene deletion. **(A)** Principal component analysis among VC and *NELL1* KO cells. **(B)** Volcano plot of 453 proteins expressed among VC and *NELL1* KO cells. Red dots indicate >2SD increase among *NELL1* KO cells. Blue dots indicate >2SD decrease among *NELL1* KO cells. **(C)** Shared GO terms enriched in both transcriptomic and proteomic analysis in *NELL1* KO cells. n=4 biological replicates of VC and *NELL1* KO cells.

**Fig. S7.**
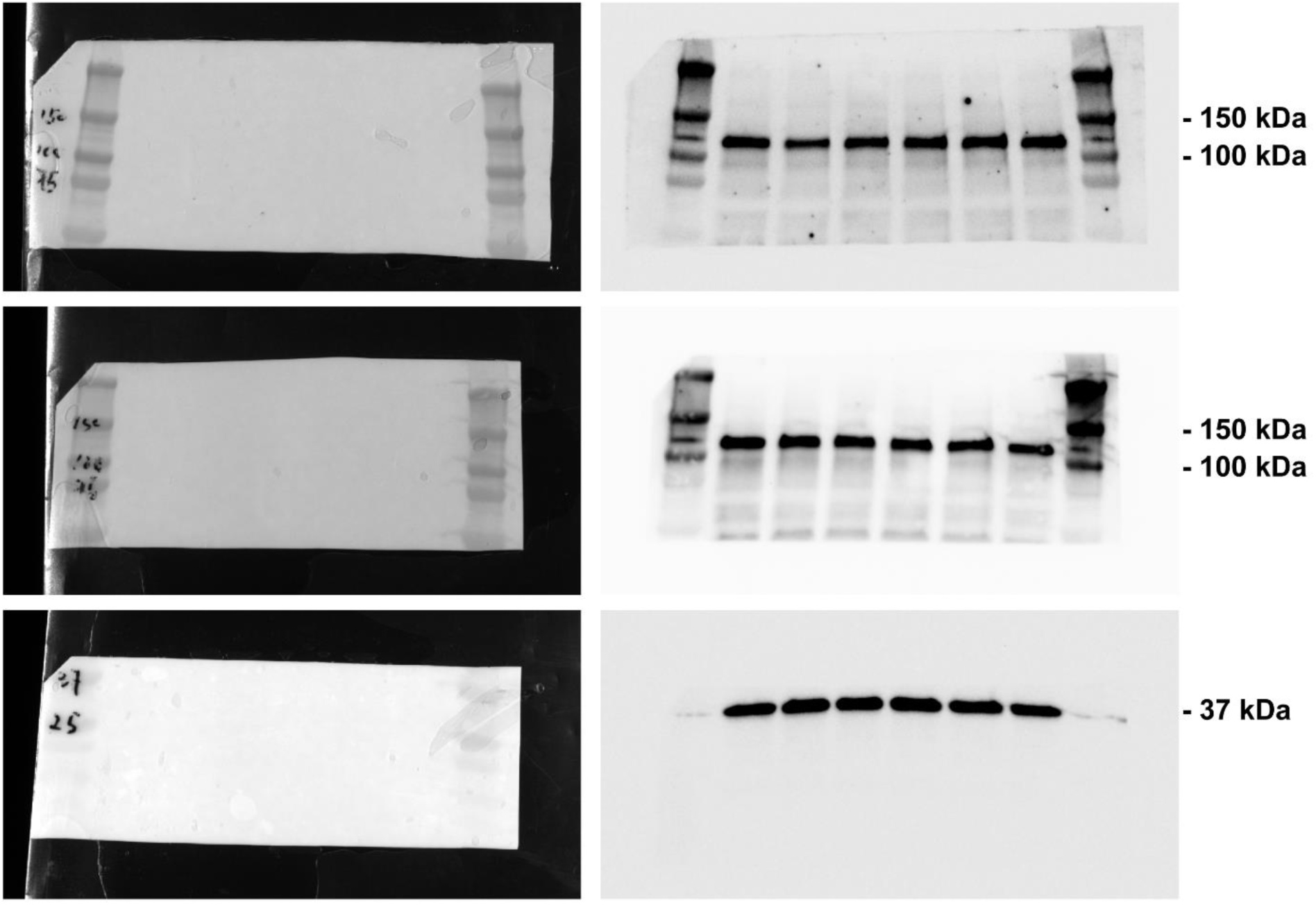
Full-size blots of Figure 5I.

**Fig. S8.**
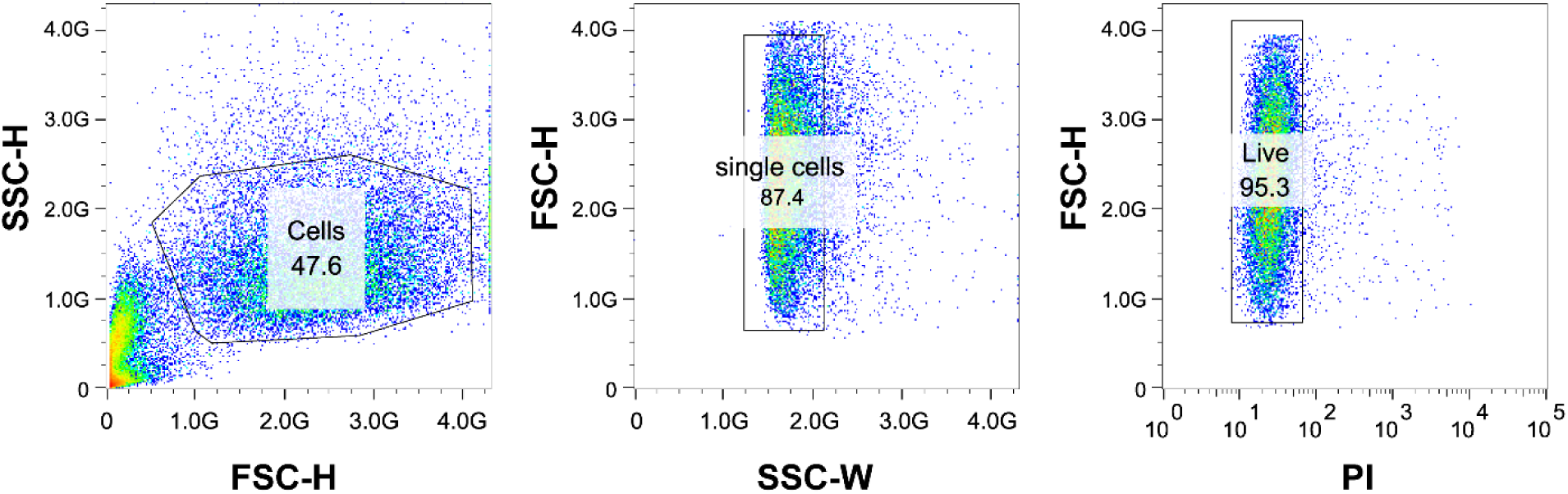
Representative flow cytometry of primary OS cells used in Figure 6C.

**Supplementary Table S1.** Tumor penetrance for mouse cell implantation model.

| Cell treatment | Total animals (N) | Animals with tumors (N) | Penetrance | Mean latency in days ± 1 SD (range) |
| --- | --- | --- | --- | --- |
| *p53<sup>fl/fl</sup>;Rb<sup>fl/fl</sup>;Cre<sup>+</sup>* (DKO) mouse BMSC | 12 | 12 | 100% | 135.2 ± 29 (85-168) |
| *p53<sup>fl/fl</sup>;Rb<sup>fl/fl</sup>;Nell1<sup>fl/fl</sup>;Cre<sup>+</sup>* (TKO) mouse BMSC | 12 | 3 | 25% | 168.0 ± 34.8 (145-208) |

**Supplementary Table S2.** Summary of tumor formation within spontaneous OS mouse model with or without *Nell1* gene deletion.

| OC-Cre | Genotype |  |  | Total animals (N) | Animals with tumors (N) | Sex distribution |  | Penetrance | Mean latency in days<br>± 1 SD<br>(range) | p-value<br>(compared with<br>corresponding<br>*Nell1*<sup>+/+</sup>) |
| --- | --- | --- | --- | --- | --- | --- | --- | --- | --- | --- |
|  | *p53* | *Rb* | *Nell1* |  |  | Male (N) | Female (N) |  |  |  |
| + | fl/fl | fl/fl | +/+ | 21 | 11 | 4 | 7 | 52.4% | 146.5 ± 28.9<br>(93-179) |  |
| + | fl/fl | fl/+ | +/+ | 21 | 10 | 6 | 4 | 47.6% | 225.3 ± 47.1<br>(112-275) |  |
| + | fl/fl | fl/fl | fl/fl | 19 | 3 | 1 | 2 | 15.8% | 398 ± 62.8<br>(339-464) | <0.0001 |
| + | fl/fl | fl/+ | fl/fl | 18 | 2 | 1 | 1 | 11.1% | 344.5 ± 19.1<br>(331-358) | 0.0066 |

**Supplementary Table S3.** Summary of spontaneous OS tumors location and incidence.

| OC-Cre | Genotype |  |  | Location | Number (Ratio) |
| --- | --- | --- | --- | --- | --- |
|  | *p53* | *Rb* | *Nell1* |  |  |
| + | fl/fl | fl/fl | +/+ | Rib/vertebra | 1 (9.1%) |
|  |  |  |  | Forelimb | 3 (27.3%) |
|  |  |  |  | Hindlimb | 7 (63.6%) |
| + | fl/fl | fl/fl | fl/fl | Forelimb | 2 (66.7%) |
|  |  |  |  | Hindlimb | 1 (33.3%) |

**Supplementary Table S4:** Most downregulated pathways among *NELL1* KO clonal 143B cells by Ingenuity Pathway Analysis.

| Pathway | Z score | -log p-value |
| --- | --- | --- |
| Dendritic Cell Maturation | -2.84 | 4.4 |
| ILK Signaling | -2.646 | 0.955 |
| ERK/MAPK Signaling | -2.646 | 0.629 |
| Coronavirus Pathogenesis Pathway | -2.53 | 2.33 |
| GP6 Signaling Pathway | -2.496 | 4.98 |
| Production of Nitric Oxide and Reactive Oxygen Species in Macrophages | -2.449 | 0.455 |
| p38 MAPK Signaling | -2.333 | 2.43 |
| IL-6 Signaling | -2.333 | 2.26 |
| Acute Phase Response Signaling | -2.333 | 2.15 |
| NF-κB Signaling | -2.333 | 1.4 |
| Protein Kinase A Signaling | -2.309 | 0.879 |
| TREM1 Signaling | -2.236 | 1.45 |
| Xenobiotic Metabolism AHR Signaling Pathway | -2.236 | 1.32 |
| Necroptosis Signaling Pathway | -2.236 | 0.44 |
| Type I Diabetes Mellitus Signaling | -2.121 | 4.69 |
| Colorectal Cancer Metastasis Signaling | -2 | 3.58 |
| Role of Pattern Recognition Receptors in Recognition of Bacteria and Viruses | -2 | 1.88 |
| Role of IL-17F in Allergic Inflammatory Airway Diseases | -2 | 1.67 |
| FAT10 Cancer Signaling Pathway | -2 | 1.5 |
| NF-κB Activation by Viruses | -2 | 0.783 |
| VEGF Family Ligand-Receptor Interactions | -2 | 0.754 |
| PI3K Signaling in B Lymphocytes | -2 | 0.567 |
| Type II Diabetes Mellitus Signaling | -2 | 0.521 |
| Tec Kinase Signaling | -2 | 0.399 |
| Integrin Signaling | -2 | 0.353 |

**Supplementary Table S5:** Most upregulated pathways among *NELL1* KO clonal 143B cells by Ingenuity Pathway Analysis.

| Pathway | Z score | -log p-value |
| --- | --- | --- |
| PPAR Signaling | 2.236 | 0.907 |
| LXR/RXR Activation | 2.121 | 1.86 |
| Mevalonate Pathway I | 2 | 3.57 |
| Superpathway of Geranylgeranyldiphosphate Biosynthesis I (via Mevalonate) | 2 | 3.08 |
| Superpathway of Cholesterol Biosynthesis | 2 | 2.25 |
| PPARα/RXRα Activation | 2 | 0.305 |
| T Cell Exhaustion Signaling Pathway | 1.633 | 1.17 |
| White Adipose Tissue Browning Pathway | 1.342 | 0.618 |

**Supplementary Table S6:** Osteosarcoma related genes in clonal 143B cells (*NELL1* KO vs. Vector control)

| Genes | Log2 FC | p-value |
| --- | --- | --- |
| L1CAM | -1.2522 | 0.00801133 |
| BIRC7 | -1.00944 | 0.196184 |
| TNFRSF11B | -0.887322 | 0.0864929 |
| VEGFA | -0.794242 | 0.0130133 |
| MMP13 | -0.729936 | 0.107346 |
| GAS7 | -0.675518 | 0.215202 |
| IER3 | -0.64576 | 0.126576 |
| MYCN | -0.56198 | 0.665071 |
| S100A4 | -0.56186 | 0.0114564 |
| BMP2 | -0.491486 | 0.313049 |
| PMP22 | -0.47866 | 3.75E-05 |
| ERRFI1 | -0.405877 | 0.384411 |
| TSPAN31 | -0.383682 | 0.0716137 |
| CASP3 | -0.35849 | 0.00365317 |
| CTSL | -0.2543 | 0.073668 |
| SRC | -0.242787 | 0.315167 |
| EXT1 | -0.220115 | 0.00830889 |
| TP53 | -0.186245 | 0.00493618 |
| KDM4C | -0.181524 | 0.419078 |
| WNT10B | -0.16363 | 0.61923 |
| HES1 | -0.10643 | 0.869479 |
| CD44 | -0.101047 | 0.669849 |
| TERF2 | -0.09173 | 0.621637 |
| STAT3 | -0.07681 | 0.446684 |
| BCL2L2 | -0.073435 | 0.531386 |
| PLK1 | -0.0622225 | 0.622098 |
| ABCG2 | -0.0363813 | 0.719462 |
| CCNG1 | -0.03209 | 0.771689 |
| ERBB2 | -0.0232025 | 0.827757 |
| BAX | 0.0027575 | 0.97162 |
| CHEK2 | 0.0217075 | 0.861121 |
| MDM2 | 0.0256225 | 0.738639 |
| MMP2 | 0.0502575 | 0.931319 |
| RB1 | 0.0632125 | 0.519254 |
| CDK4 | 0.121727 | 0.0655916 |
| CDKN2D | 0.186561 | 0.364097 |
| RECQL4 | 0.18896 | 0.343264 |
| MCM4 | 0.21399 | 0.0494394 |
| FOS | 0.21461 | 0.879092 |
| PTH1R | 0.216413 | 0.292116 |

| **Genes** | **Log2 FC** | **p-value** |
| --- | --- | --- |
| PRIM1 | 0.27913 | 5.03E-05 |
| CCNB2 | 0.280205 | 0.100503 |
| KIT | 0.310382 | 0.157051 |
| HMGB1 | 0.363268 | 0.00364131 |
| TNFRSF11A | 0.380036 | 0.588856 |
| BAD | 0.382345 | 0.0062038 |
| RMI1 | 0.487102 | 0.00328483 |
| CCNE1 | 0.553267 | 0.00877644 |
| CCN1 | 0.61587 | 0.184948 |
| SPARC | 0.620737 | 0.00929151 |
| E2F2 | 0.886565 | 0.00102943 |
| BCL2 | 0.950508 | 0.004066 |

**Supplementary Table S7:** Most downregulated Gene Ontology terms among *NELL1* KO clonal 143B cells by RNA sequencing.

| Gene Ontology | Count | p-value |
| --- | --- | --- |
| GO:0030198 extracellular matrix organization | 50 | 1.28E-17 |
| GO:0001525 angiogenesis | 41 | 9.90E-10 |
| GO:0030574 collagen catabolic process | 20 | 5.74E-09 |
| GO:0007155 cell adhesion | 62 | 1.26E-08 |
| GO:0035987 endodermal cell differentiation | 13 | 2.18E-08 |
| GO:0030334 regulation of cell migration | 18 | 2.03E-06 |
| GO:0007229 integrin-mediated signaling pathway | 21 | 2.28E-06 |
| GO:0070059 intrinsic apoptotic signaling pathway in response to endoplasmic reticulum stress | 11 | 1.98E-05 |
| GO:0042981 regulation of apoptotic process | 31 | 2.09E-05 |
| GO:0010575 positive regulation of vascular endothelial growth factor production | 10 | 2.20E-05 |
| GO:0007165 signal transduction | 106 | 4.76E-05 |
| GO:0022617 extracellular matrix disassembly | 16 | 5.48E-05 |
| GO:0042127 regulation of cell proliferation | 27 | 7.52E-05 |
| GO:0071260 cellular response to mechanical stimulus | 15 | 9.74E-05 |
| GO:0008285 negative regulation of cell proliferation | 45 | 1.20E-04 |
| GO:0043123 positive regulation of I-kappaB kinase/NF-kappaB signaling | 24 | 1.49E-04 |
| GO:0001666 response to hypoxia | 25 | 1.57E-04 |
| GO:0051092 positive regulation of NF-kappaB transcription factor activity | 21 | 1.96E-04 |
| GO:0048333 mesodermal cell differentiation | 6 | 3.04E-04 |
| GO:0043066 negative regulation of apoptotic process | 48 | 3.96E-04 |
| GO:0001570 vasculogenesis | 12 | 5.53E-04 |
| GO:0045766 positive regulation of angiogenesis | 18 | 7.04E-04 |
| GO:0045599 negative regulation of fat cell differentiation | 10 | 9.02E-04 |
| GO:0043065 positive regulation of apoptotic process | 34 | 9.72E-04 |
| GO:0006915 apoptotic process | 55 | 0.001069105 |

**Supplementary Table S8:** Most upregulated Gene Ontology terms among *NELL1* KO clonal 143B cells by RNA sequencing.

| Gene Ontology | Count | p-value |
| --- | --- | --- |
| GO:0006260 DNA replication | 34 | 6.06E-11 |
| GO:0006695 cholesterol biosynthetic process | 16 | 1.17E-09 |
| GO:0000732 strand displacement | 12 | 8.42E-08 |
| GO:0006281 DNA repair | 37 | 9.26E-08 |
| GO:0000731 DNA synthesis involved in DNA repair | 13 | 3.37E-07 |
| GO:0036297 interstrand cross-link repair | 13 | 1.75E-05 |
| GO:0007067 mitotic nuclear division | 32 | 4.86E-05 |
| GO:0051301 cell division | 40 | 7.51E-05 |
| GO:0008299 isoprenoid biosynthetic process | 7 | 7.66E-05 |
| GO:0000082 G1/S transition of mitotic cell cycle | 18 | 7.74E-05 |
| GO:0007059 chromosome segregation | 14 | 1.28E-04 |
| GO:0000083 regulation of transcription involved in G1/S transition of mitotic cell cycle | 8 | 2.38E-04 |
| GO:0055114 oxidation-reduction process | 56 | 3.94E-04 |
| GO:0030324 lung development | 14 | 4.04E-04 |
| GO:0006974 cellular response to DNA damage stimulus | 26 | 4.61E-04 |
| GO:0003139 secondary heart field specification | 5 | 6.60E-04 |
| GO:0034080 CENP-A containing nucleosome assembly | 10 | 6.91E-04 |
| GO:0008283 cell proliferation | 38 | 7.65E-04 |
| GO:0006099 tricarboxylic acid cycle | 8 | 0.001113854 |
| GO:0000122 negative regulation of transcription from RNA polymerase II promoter | 63 | 0.001240728 |
| GO:0001568 blood vessel development | 9 | 0.001307687 |
| GO:0035264 multicellular organism growth | 13 | 0.00217589 |
| GO:0010628 positive regulation of gene expression | 28 | 0.002846398 |
| GO:0060394 negative regulation of pathway-restricted SMAD protein phosphorylation | 5 | 0.003868161 |
| GO:0045746 negative regulation of Notch signaling pathway | 7 | 0.005722524 |

**Supplementary Table S9:** Most downregulated Gene Ontology terms among *NELL1* KO clonal 143B cells by proteomic analysis.

| Gene Ontology | Count | p-value |
| --- | --- | --- |
| GO:0043066 negative regulation of apoptotic process | 13 | 3.17E-08 |
| GO:0008284 positive regulation of cell proliferation | 13 | 4.13E-08 |
| GO:0000082 G1/S transition of mitotic cell cycle | 8 | 5.11E-08 |
| GO:0046777 protein autophosphorylation | 9 | 1.11E-07 |
| GO:0051726 regulation of cell cycle | 8 | 1.97E-07 |
| GO:0042127 regulation of cell proliferation | 8 | 2.92E-06 |
| GO:0008283 cell proliferation | 10 | 3.60E-06 |
| GO:0018105 peptidyl-serine phosphorylation | 7 | 4.01E-06 |
| GO:0042493 response to drug | 9 | 8.05E-06 |
| GO:0032355 response to estradiol | 6 | 1.40E-05 |
| GO:0008285 negative regulation of cell proliferation | 9 | 5.36E-05 |
| GO:0000083 regulation of transcription involved in G1/S transition of mitotic cell cycle | 4 | 6.26E-05 |
| GO:0000122 negative regulation of transcription from RNA polymerase II promoter | 11 | 1.37E-04 |
| GO:0006281 DNA repair | 7 | 1.41E-04 |
| GO:0035264 multicellular organism growth | 5 | 1.56E-04 |
| GO:0018108 peptidyl-tyrosine phosphorylation | 6 | 1.68E-04 |
| GO:0009636 response to toxic substance | 5 | 1.97E-04 |
| GO:0007169 transmembrane receptor protein tyrosine kinase signaling pathway | 5 | 3.14E-04 |
| GO:0006469 negative regulation of protein kinase activity | 5 | 3.53E-04 |
| GO:0035556 intracellular signal transduction | 8 | 4.09E-04 |
| GO:0045471 response to ethanol | 5 | 4.42E-04 |
| GO:0048015 phosphatidylinositol-mediated signaling | 5 | 4.58E-04 |
| GO:0030335 positive regulation of cell migration | 5 | 0.003505609 |
| GO:0045747 positive regulation of Notch signaling pathway | 3 | 0.005587098 |
| GO:0034446 substrate adhesion-dependent cell spreading | 3 | 0.007358625 |

**Supplementary Table S10:** Most upregulated Gene Ontology terms among *NELL1* KO clonal 143B cells by proteomic analysis.

| Gene Ontology | Count | p-value |
| --- | --- | --- |
| GO:0043065 positive regulation of apoptotic process | 8 | 3.54E-07 |
| GO:0071260 cellular response to mechanical stimulus | 5 | 4.78E-06 |
| GO:0071456 cellular response to hypoxia | 5 | 1.59E-05 |
| GO:0048661 positive regulation of smooth muscle cell proliferation | 4 | 1.19E-04 |
| GO:0001666 response to hypoxia | 5 | 1.55E-04 |
| GO:0006915 apoptotic process | 7 | 2.33E-04 |
| GO:0042981 regulation of apoptotic process | 5 | 3.51E-04 |
| GO:0008637 apoptotic mitochondrial changes | 3 | 4.19E-04 |
| GO:0043066 negative regulation of apoptotic process | 6 | 7.04E-04 |
| GO:0045944 positive regulation of transcription from RNA polymerase II promoter | 8 | 7.19E-04 |
| GO:0098609 cell-cell adhesion | 5 | 8.69E-04 |
| GO:0097193 intrinsic apoptotic signaling pathway | 3 | 0.001053376 |
| GO:1900740 positive regulation of protein insertion into mitochondrial membrane involved in apoptotic signaling pathway | 3 | 0.001053376 |
| GO:0042493 response to drug | 5 | 0.001330423 |
| GO:0008630 intrinsic apoptotic signaling pathway in response to DNA damage | 3 | 0.002574001 |
| GO:0042542 response to hydrogen peroxide | 3 | 0.003023949 |
| GO:0008285 negative regulation of cell proliferation | 5 | 0.003479001 |
| GO:0001525 angiogenesis | 4 | 0.005342541 |
| GO:0007584 response to nutrient | 3 | 0.006262053 |
| GO:0032869 cellular response to insulin stimulus | 3 | 0.006763636 |
| GO:0030307 positive regulation of cell growth | 3 | 0.008002639 |
| GO:0072656 maintenance of protein location in mitochondrion | 2 | 0.008014682 |
| GO:0072655 establishment of protein localization to mitochondrion | 2 | 0.009610179 |
| GO:0032287 peripheral nervous system myelin maintenance | 2 | 0.011203204 |
| GO:0016032 viral process | 4 | 0.011917614 |

**Supplementary Table S11.** Summary of NELL1 and FN1 immunostaining in biopsy samples of high grade conventional osteosarcoma.

| Sample | Biopsy (%Area) |  |
| --- | --- | --- |
|  | NELL1 | FN1 |
| #1 | 38.6 | 50.5 |
| #2 | 28.2 | 38.7 |
| #3 | 23.6 | 18.4 |
| #4 | 18.1 | 21.1 |
| #5 | 13.9 | 11.0 |

**Supplementary Table S12.** Antibodies used.

| **Antibody** | **Company** | **Catalog #** | **Use** |
| --- | --- | --- | --- |
| Rabbit anti-CD31 | Abcam | ab28364 | IF |
| Rat anti-Mouse CD31 | BD Pharmingen | 551262 | F |
| Mouse anti-Human CD44 | BD Pharmingen | 561289 | F |
| Rat anti-Mouse CD45 | BD Pharmingen | 559864 | F |
| Mouse anti-Human CD73 | BD Pharmingen | 561014 | F |
| Mouse anti-Human CD90 | BD Pharmingen | 555595 | F |
| Mouse anti-Human CD105 | BD Pharmingen | 562380 | F |
| Rabbit anti-FAK | Abcam | ab40794 | IF/WB |
| Rabbit anti-pFAK | Abcam | ab81298 | IF/WB/ICC |
| Rabbit anti-Fibronectin | Abcam | ab2413 | IF |
| Rabbit anti-GAPDH | Abcam | ab9485 | WB |
| Rabbit anti-Ki67 | Abcam | ab15580 | IF |
| Mouse anti-Laminin | Cell signaling | 4777S | IF |
| Mouse anti-human NELL1 | R&D system | MAB54871 | ICC/IF |
| Anti-rabbit IgG, HRP-linked Antibody | Cell Signaling Technology | 7074 | WB |
| Goat anti-Mouse AF488 | Abcam | ab150117 | IF |
| Goat anti-Rabbit AF488 | Abcam | ab150077 | IF |
| Goat anti-Rabbit DyLight 594 | Vector Laboratories | DI-1594 | IF |
| Donkey anti-Goat AF647 | Abcam | ab150135 | IF |
| Goat anti-Mouse AF647 | Abcam | ab150119 | IF |
| Goat anti-Rabbit AF647 | Abcam | ab150079 | IF |
| Goat anti-Rat AF647 | Abcam | ab150167 | IF |
| F: Flow cytometry; IF: Immunofluorescent staining; IHC: Immunohistochemistry; ICC: Immunocytochemistry. |  |  |  |

**Supplementary Table S13.**
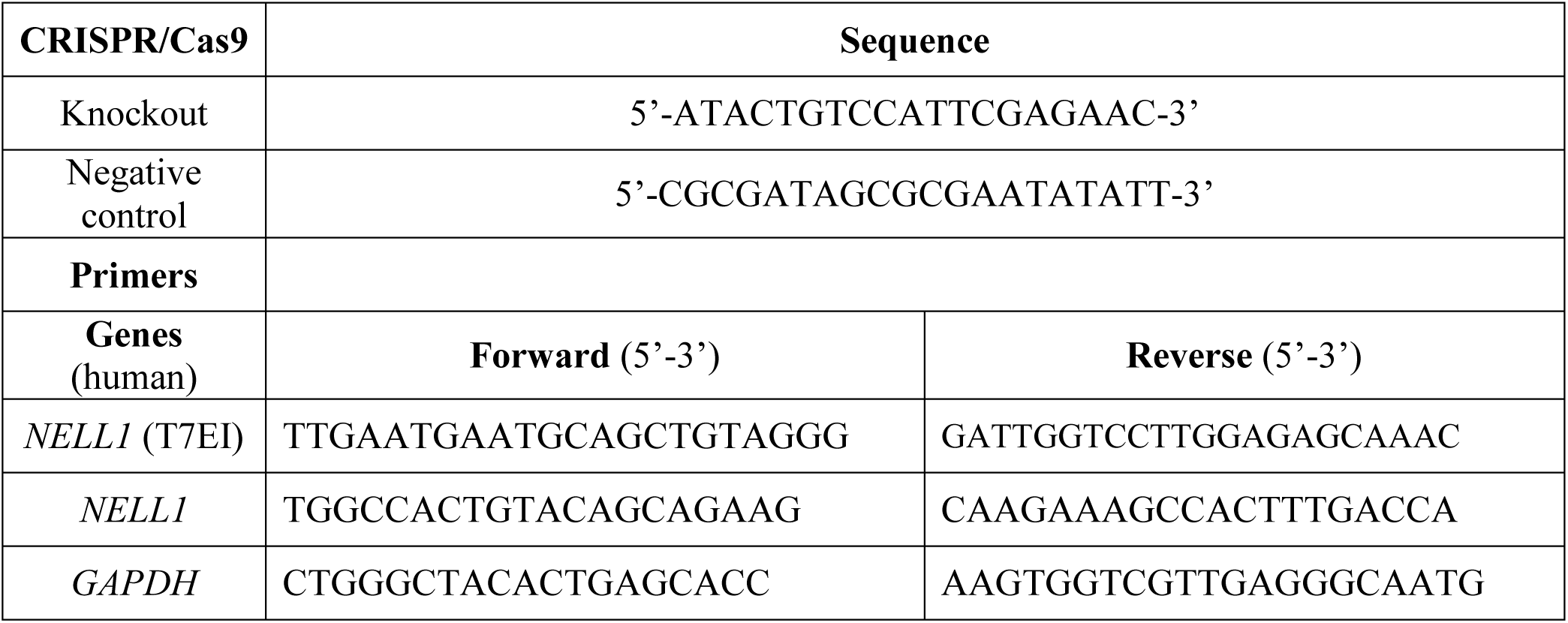
CRISPR/Cas9 sgRNA and primer sequences.

**Supplementary Table S14.** Summary of osteosarcoma cell implantation models.

| Study | Cell type | Injection location | Cell # per animal | # of animals | Endpoint |
| --- | --- | --- | --- | --- | --- |
| Tumor size | Vector control 143B | Tibial periosteum | 1 M | 8 | 28 d |
|  | *Nell1* KO 143B, clone #2 | Tibial periosteum | 1 M | 8 | 28 d |
| Overall survival | Vector control 143B | Tibial periosteum | 1 M | 10 | Until moribund |
|  | *Nell1* KO 143B, clone #2 | Tibial periosteum | 1 M | 10 | Until moribund |
| Pulmonary metastasis | Vector control 143B | Intravenous (tail vein) | 1.5 M | 8 | 28 d |
|  | *Nell1* KO 143B, clone #2 | Intravenous (tail vein) | 1.5 M | 8 | 28 d |
| Tumor size | Vector control 143B | Tibial periosteum | 1 M | 9 | 28 d |
|  | *Nell1* KO 143B, clone #4 | Tibial periosteum | 1 M | 13 | 28 d |
| Tumor free survival | *p53*<sup>fl/fl</sup>;*Rb*<sup>fl/fl</sup>;Cre<sup>+</sup> (DKO) mouse BMSC | Tibial periosteum | 1 M | 12 | 28 d after tumor observed |
|  | *p53*<sup>fl/fl</sup>;*Rb*<sup>fl/fl</sup>;*Nell1*<sup>fl/fl</sup>;Cre<sup>+</sup> (TKO) mouse BMSC | Tibial periosteum | 1 M | 12 | 28 d after tumor observed |
| For survival studies, mice were monitored three times weekly. |  |  |  |  |  |

## Data availability

RNA sequencing data is freely available within the NCBI GEO database (GSE182703). Proteomic data is submitted as a separate Excel file.

